# A common algorithm for confidence judgements across visual, auditory and audio-visual decisions

**DOI:** 10.1101/2025.02.12.637846

**Authors:** Rebecca K West, Natasha Matthews, Jason B Mattingley, David K Sewell

## Abstract

Most studies investigating the computational basis of decision confidence have focused on simple visual perceptual tasks, leaving open questions about how confidence is formed in decisions involving other sensory modalities or those requiring the integration of information across modalities. To address these gaps, we used computational modelling to analyse confidence judgements in perceptual decisions involving visual, auditory, and audio-visual stimuli. Drawing on research into visual confidence, we adapted models from the literature to evaluate their fit to our data, comparing three popular classes: unscaled evidence strength, scaled evidence strength, and Bayesian models. Our results show that the scaled evidence strength models consistently outperformed the other model classes across all tasks and could also be used to predict behaviour in the audio-visual task from the unidimensional auditory and visual model fits. These findings suggest that confidence judgements across different perceptual decisions rely on a shared algorithm that dynamically accounts for both sensory uncertainty and evidence strength, without the computation of posterior probabilities. Additionally, we investigated the algorithms used for multidimensional (audio-visual) confidence judgements specifically, showing that participants integrated both the visual and auditory dimensions of the stimulus, rather than relying solely on the most informative modality, and used a modality-independent measure of sensory uncertainty to adjust their confidence. Overall, our findings provide evidence for a common algorithm underlying confidence judgements across modalities and demonstrate the broad applicability of the scaled evidence strength algorithm, even in tasks requiring the integration of distinct sensory information.

## Introduction

Human perceptual decisions are often accompanied by a level of confidence that reflects the accuracy of the decision (Boldt et al., 2017; Kepecs et al., 2008). This ability to accurately monitor and evaluate one’s decision-making processes, referred to as *metacognition*, plays a crucial role in humans’ capacity to make adaptive decisions. It prompts individuals to seek more information before committing to a choice (Desender et al., 2018), decide when to stop deliberating in a difficult decision (van den Berg et al., 2016), or anticipate effort or uncertainty in an upcoming decision (Boldt et al., 2019). Given the pivotal role that metacognitive judgements play in adaptive decision-making, one crucial goal for cognitive science has been to understand the psychological processes involved in generating such judgements.

Investigating the mechanisms that underpin metacognitive judgements, however, requires considering how these computations are shaped by the sensory modalities involved. For instance, forming confidence about the colour of a traffic light relies solely on visual information, confidence in whether a phone is ringing relies on auditory input, and confidence in whether a pineapple is ripe requires integrating information about its colour, smell, and texture (de Gardelle et al., 2016). Most previous studies investigating confidence judgements, however, focused on simple visual perceptual decisions only (Adler & Ma, 2018; Aitchison et al., 2015; Denison et al., 2018; Hangya et al., 2016; Li & Ma, 2020; Lisi et al., 2021; Locke et al., 2022; Navajas et al., 2017; Sanders et al., 2016).

To bridge this significant gap in our understanding about the formation of confidence across sensory modalities, our previous work utilised computational modelling to explore the psychological algorithms underlying confidence judgements across different perceptual domains (West et al., 2023). Specifically, we investigated whether the algorithms underlying confidence judgements were the same across vision and audition. Participants completed two versions of a categorisation task with visual and auditory stimuli, where they categorised each stimulus based on a perceptual feature (orientation or frequency) and then reported their confidence in that decision. In both modalities, we varied evidence strength (i.e., the strength of the evidence for a particular category) and sensory uncertainty (i.e. the amount of noise or uncertainty in the sensory signal). This allowed us to keep the decision context the same but vary the modality of presentation, isolating any similarities (*modality-independent* processes) or differences (*modality-specific* processes) in the algorithms used to generate confidence. In order to quantify the algorithms used for confidence, we evaluated the performance of several classes of existing models which mathematically formalise the mapping of evidence strength and sensory uncertainty to confidence in different ways. We found that a single class of models, in which confidence is derived from a subjective assessment of the strength of the evidence for a particular choice, scaled by an estimate of sensory uncertainty, provided the best account of confidence for both visual and auditory decisions. Comparing the ‘settings’ (or parameters) of this process across modalities however, revealed that the algorithm was tuned differently in each modality. That is, we found evidence for a modality-general scaled evidence strength algorithm that was tuned specifically in each modality (Lehmann et al., 2022; Masset et al., 2020; Mazancieux et al., 2020; Morales et al., 2018; Rouault et al., 2018).

While our recent work (West et al., 2023) has expanded our understanding of the algorithms that underlie confidence judgements in distinct sensory modalities, many everyday decisions require the integration of information from multiple senses. The psychological computations supporting metacognitive judgements in multidimensional contexts, however, remain under-researched and poorly understood. To address this gap, the current study explored both how participants make metacognitive judgements for multidimensional perceptual decisions and how the algorithms used for multidimensional decisions compare with those used in unidimensional contexts-those involving a single sensory modality.

In exploring the processes involved in multidimensional decision-making, we can build on substantial research that has investigated how individuals combine information from different senses, such as vision and audition, to form a unified perceptual representation. Specifically, cue combination research has explored whether perceptual judgements are formed by combining inputs from different senses in a weighted manner (integration perspective) or if they are based on a maximal input approach (selection perspective), where the brain selects only the dominant sensory input. Numerous studies have shown that individuals combine information from multiple sensory modalities by weighting inputs based on their reliability, which is often referred to as optimal integration or Bayesian cue integration. According to weighted integration models, sensory input from different modalities is not treated equally; instead more reliable (less noisy) information is given greater weight when integrating cues (Ernst & Banks, 2002; McGurk & MacDonald, 1967). Other studies suggest that, in certain situations, rather than combining cues, people rely predominantly on the most informative or salient sensory modality (Posner et al., 1976; Welch & Warren, 1980). For example, under conditions of uncertainty or conflict, perceptual judgements often favour the most easily processed modality (Posner et al., 1976; Welch & Warren, 1980).

The same question of how sensory input is combined across modalities also applies to confidence judgements. Specifically, it is not known whether confidence is based on a weighted integration of information from different senses, or if it follows a selection-based approach, where confidence reflects input from the most informative or salient modality only. Recent findings from Gao et al. (2023) support the weighted integration perspective, showing that participants combined auditory and visual motion cues based on their reliability. In Gao et al.’s multisensory task, participants were instructed to judge the direction of an auditory motion cue and ignore a simultaneously presented visual motion cue. Despite these instructions, they found that participants’ judgements and confidence were influenced by both the direction and reliability of the visual cue. This effect was captured by a computational model which assumed that the cognitive processes involved in confidence judgements are the same as those used in lower-level stages of perception. These findings also align with the results of Faivre et al. (2018) who used computational modelling to demonstrate that confidence in an audio-visual task could be predicted by an integrated representation of auditory and visual signals. Specifically, Faivre and colleagues (2018) modelled confidence as the probability of being correct, showing that participants’ perceived probability of correctness depended on the integration of sensory inputs from both visual and auditory modalities.

Building on these studies, here we also investigated how confidence judgements operate in multidimensional contexts by using a task in which stimulus information provided in different modalities was explicitly task relevant (Gao et al., 2023), and by expanding the range of models used to investigate confidence beyond purely probabilistic frameworks (Faivre et al., 2018). This approach enabled us to compare multiple potential algorithms for generating confidence, offering a more nuanced comparison between the integration-based and selection-based perspectives. Moreover, by using an expanded modelling framework, we were able to explore potential biases in integration as well as more directly investigate the role of prior knowledge in confidence judgements, providing deeper insight into the cognitive processes underlying multidimensional confidence formation.

A second, related question that we addressed in this study was whether the algorithms used for confidence judgements differ between unidimensional stimuli, such as a purely visual or auditory stimulus, and multidimensional stimuli, such as an audio-visual stimulus. A modality-general framework suggests that the same computational processes apply across both uni- and multidimensional contexts, where a single algorithm is tuned to the specific decision context. However, one alternative view posits that integrating information from multiple senses may require a distinct computational strategy, possibly leading multidimensional decisions to rely on optimal or near-optimal, Bayesian-like algorithms. That is, while simpler heuristic strategies may be sufficient for unidimensional contexts (Adler & Ma, 2018; West et al., 2023), a more statistically optimal approach could be better suited for multidimensional contexts. Bayesian algorithms assume the computation of a probability metric which automatically resolves any ‘translation issues’ when combining (or comparing) sensory input from different modalities. Moreover, given the multidimensional nature of real-world experiences, along with the richer feedback opportunities they provide, optimal integration strategies may be more finely tuned for scenarios involving multiple sensory inputs. To assess whether a unified computational framework can account for both unidimensional and multidimensional judgements, we included unidimensional visual and auditory and multidimensional audio-visual decisions in our study.

To characterise and compare the computational processes used for metacognitive judgements in visual, auditory and audio-visual decisions, we use the same modelling framework as in our previous work (West et al., 2023). Specifically, we considered different theoretical models of decision confidence – Bayesian models and non-Bayesian evidence strength-based models – and evaluated their performance across the different types of perceptual decisions (see **Figure 1**). We provide a brief overview of the different model classes below.

**Figure 1.**
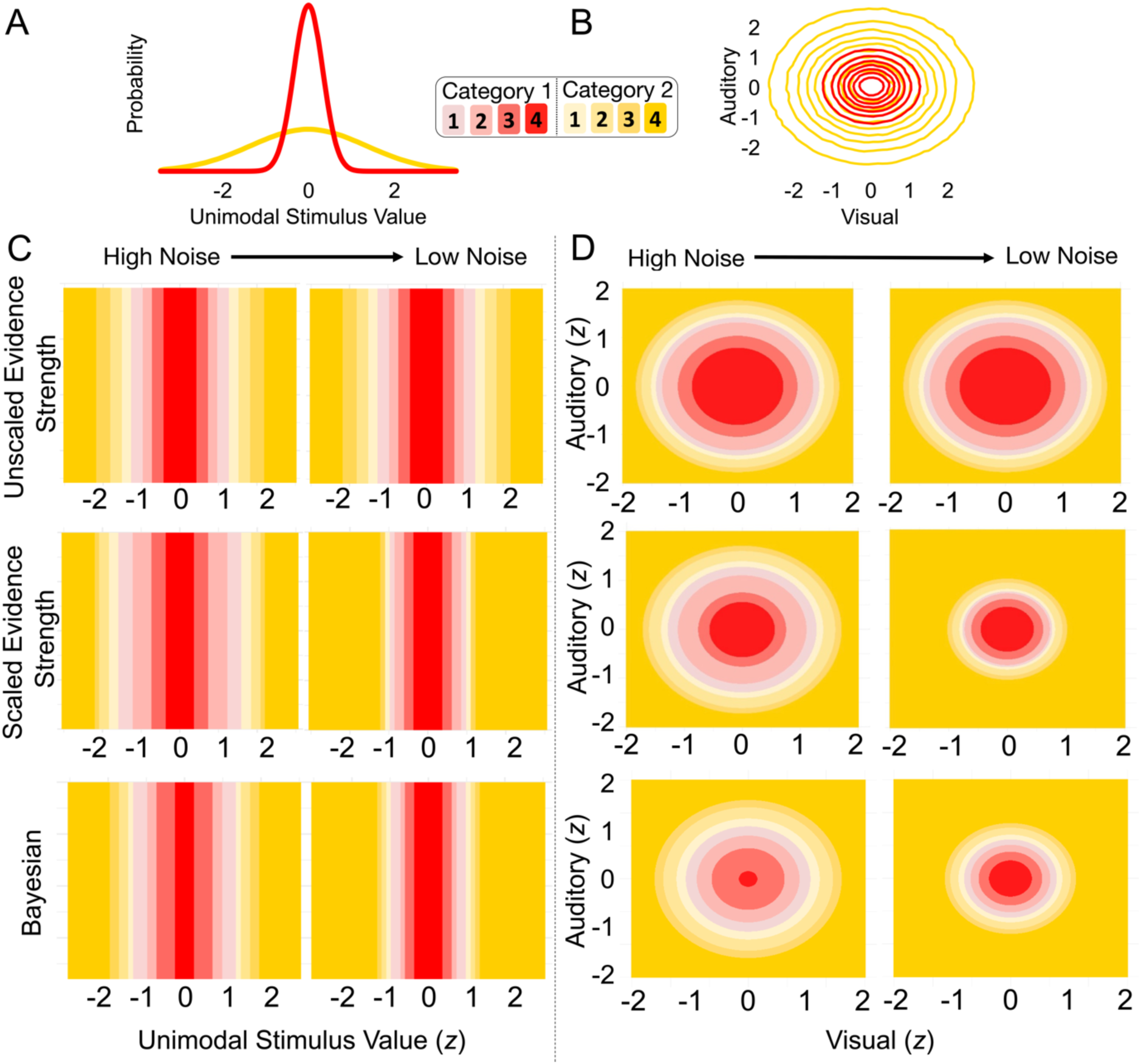
Choice and Confidence Criteria Across Models. *Note.* **(A**) Illustration of category structures for unidimensional task and **(B)** multidimensional task. For the unscaled evidence strength models (top row), the choice and confidence criteria do not change as a function of sensory noise for both the **(C)** unidimensional and **(D)** multidimensional tasks. For the scaled evidence strength models and Bayesian models (middle and bottom rows, respectively), the choice and confidence criteria change as a function of sensory uncertainty/noise.

According to *Bayesian models*, observers optimally combine knowledge about the statistical structure of a task (the “prior”) with the present sensory input (the “likelihood”) to compute a posterior probability distribution over possible states of a stimulus (Drugowitsch et al., 2014; Hangya et al., 2016; Kepecs & Mainen, 2012; Meyniel et al., 2015; Pouget et al., 2016). The posterior probability distribution thus represents every possible outcome and its associated probability. Bayesian models stipulate that the observer uses the posterior probability distribution to guide both their choice and confidence. For example, one possible strategy posits that an observer’s confidence increases with the posterior probability of the chosen outcome. In experimental paradigms where numerical confidence ratings are used, a set of criteria can map posterior probabilities to each level of the confidence rating scale. In the example of judging the ripeness of a pineapple, a Bayesian observer would eat the pineapple where *p*(*ripe* | *pineapple*) is above 0.5, with higher probabilities indicating greater confidence in the decision. Another method for mapping posterior probability to confidence is to use the log posterior probability ratio, 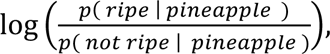 which is unbounded and allows for greater differences in confidence at different levels of sensory uncertainty (Adler & Ma, 2018; Locke et al., 2022).

*Unscaled evidence strength models* posit that confidence depends entirely on the strength of the evidence for a particular decision outcome. These models assume that the observer uses the attribute value of the stimulus to directly quantify evidence strength, with confidence increasing as evidence becomes stronger. For example, in this framework, the more saturated the yellow colour of a pineapple, the more confident the person is that it’s ripe. The relationship between evidence strength and confidence can also be quantified numerically by setting criteria based on evidence strength. Unlike Bayesian models and scaled evidence strength models (described below), unscaled evidence strength models do not explicitly account for sensory uncertainty in the computation of confidence, focusing instead on the direct relationship between attribute value and confidence.

*Scaled evidence strength models* posit that confidence depends on both the strength of the evidence and the amount of sensory uncertainty associated with the decision. According to scaled evidence strength models, sensory uncertainty is used to scale evidence strength such that under conditions of high uncertainty, greater evidence strength is required to produce the same level of confidence as under conditions of low sensory uncertainty. For example, these models suggest that a person would feel less confident about the ripeness of a pineapple whose colour is highly variable relative to one where there is low variability in colour, even though the average ‘yellowness’ of the two pineapples is the same. For numerical confidence ratings, a point estimate of sensory uncertainty can be used to scale the criteria used to delineate continuous evidence strength (Adler & Ma, 2018; Denison et al., 2018; West et al., 2023). Although both scaled evidence strength and Bayesian models consider evidence strength and sensory uncertainty as important for confidence, scaled evidence strength models do not rely on probabilistic inference. Because scaled evidence strength models do not involve the computation of a probability metric, they are considered less computationally complex, and arguably, a more feasible explanation of human cognition (Sanborn et al., 2010; Sanborn & Chater, 2016; Stewart et al., 2006).

### The Current Study

The aim of the current study was to differentiate between these different theoretical accounts and use computational models to characterise the computations involved in confidence for visual, auditory and audio-visual decisions. Specifically, one key focus of our analyses was to investigate whether the same computational algorithms governed confidence judgements across all three types of decisions. To address this question, we evaluated the performance of the different model classes and tested whether a single class consistently fit the data across tasks. We also examined the generalisability of the models by determining how well predictions from the unidimensional tasks (visual and auditory) align with observed behaviour in the audio-visual task. A second key focus of our analyses was to understand how participants integrate sensory information from multiple modalities when making multidimensional confidence judgements. To characterise the algorithms supporting these judgements, we systematically explored different assumptions regarding how sensory inputs are combined across modalities within each model class.

## Method

### Overview

Participants completed 3 categorisation tasks: (1) a visual categorisation task, (2) an auditory categorisation task and (3) an audio-visual categorisation task. In each task, participants made a two-alternative forced choice category decision (Category 1 or Category 2), followed by a 4-point confidence rating, ranging from low confidence (1) to high confidence (4). Category membership was determined by an attribute of the stimulus: orientation for visual stimuli (drifting Gabor patches), frequency for auditory stimuli (pure tones) and both orientation and frequency for the audio-visual stimuli (drifting Gabor patches with simultaneously presented pure tones). A normal distribution of the relevant stimulus attribute defined each category where the category distributions had the same mean and different standard deviations. Sensory uncertainty also varied with stimuli presented at 4 different intensities. In the visual tasks, intensity depended on the contrast of the Gabor patch (3.3%, 5%, 6.7%, 13.5%). In the auditory tasks, intensity depended on the loudness of the tone (equivalent of -3 dB SPL, 4 dB SPL, 10.5 dB SPL, 17 dB SPL for a 1 kHz pure tone). In the audio-visual task, both contrast (3.3%, 4.2%, 5.3%, 6.7%) and loudness (equivalent of -3 dB SPL, 2 dB SPL, 6 dB SPL, 10.5 dB SPL for a 1 kHz pure tone) varied together. The intensity levels were chosen to achieve a similar range of performance across all tasks based on pilot data.

### Participants

Participants were recruited from the University of Queensland’s research participation scheme and were reimbursed for their time ($20 per hour in cash or gift cards). Inclusion criteria required normal (or corrected-to-normal) vision and normal hearing, both assessed by self-report. After exclusions (see the Results section for details), the final sample comprised of 10 participants.

### Categorisation Task

As shown in **Figure 2A**, the categories were defined by normal distributions which had the same mean and different standard deviations. In the visual modality, larger rotations clockwise or counter clockwise relative to horizontal were more likely to be sampled from category 2 (*μ*_*cat*2_ = 0°, *σ*_*cat*2_ = 12°) whereas smaller rotations relative to horizontal were more likely to be sampled from category 1 (*μ*_*cat*1_ = 0°, *σ*_*cat*1_ = 3°). In the auditory modality, lower and higher frequency tones were more likely to be sampled from category 2 (*μ*_*cat*2_ = 2700 *Hz*, *σ*_*cat*2_ = 500 *Hz*), whereas intermediate frequency tones were more likely to be sampled from category 1 (*μ*_*cat*1_ = 2700 *Hz*, *σ*_*cat*1_ = 125 *Hz*). In the audio-visual task, the categories were defined by bivariate normal distributions with the same means and standard deviations as in the visual and auditory versions of the tasks (*μ*_*cat*1_ = [0°, 2700 *Hz*]; *σ*_*cat*1_ = [3°, 125 *Hz*]; *μ*_*cat*2_ = [0°, 2700 *Hz*]; *σ*_*cat*2_ = [12°, 500 *Hz*]). To maximise correct responses in each task, the ideal observer would use the points where the category distributions intersect, reporting that stimulus values contained within the intervals were sampled from category 1 and stimulus values outside the intervals were sampled from category 2 (see dotted lines in **Figure 2A**).

**Figure 2.**
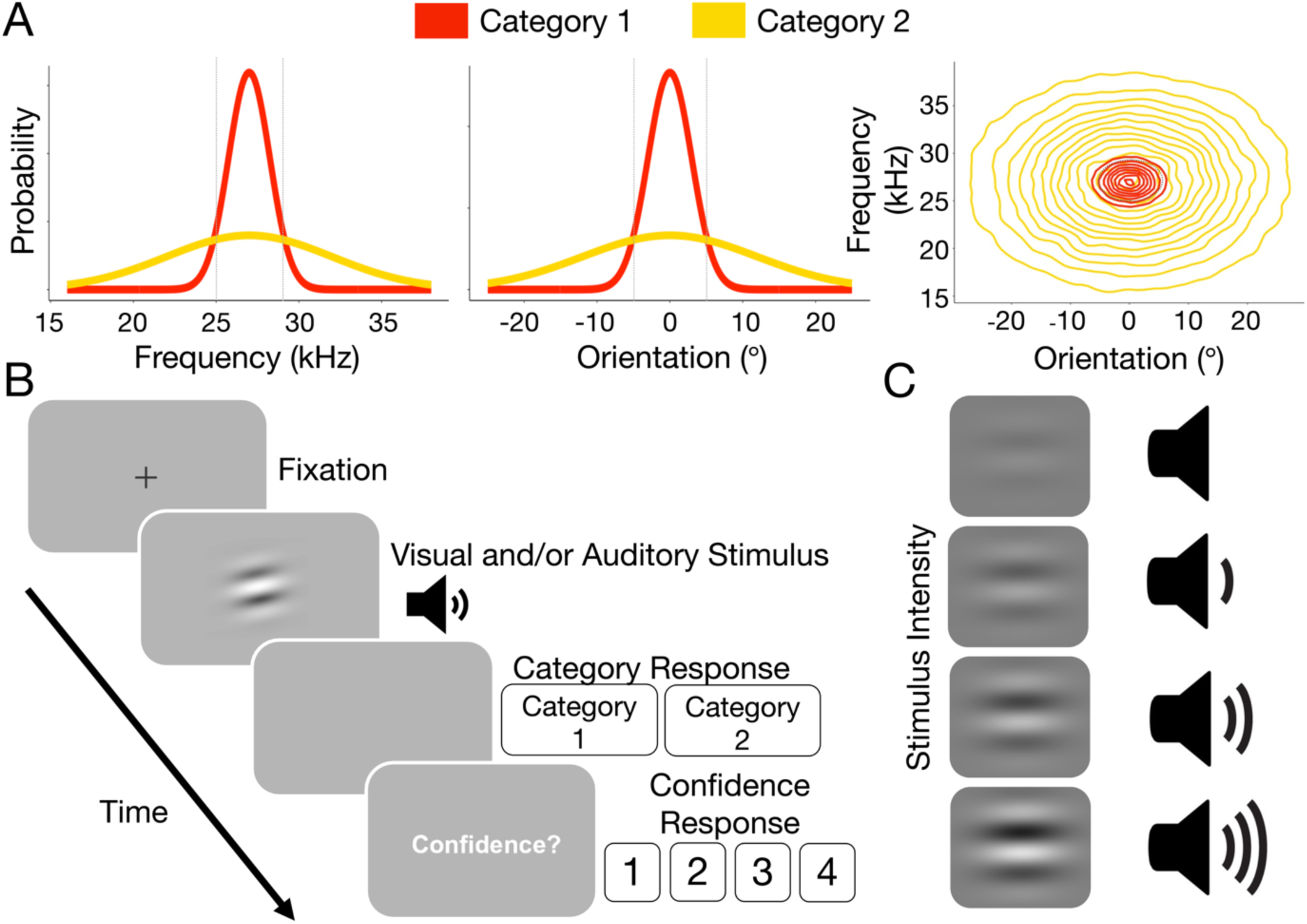
Experimental Task. *Note.* **(A)** Category distribution for auditory (left), visual (middle) and audio-visual (right) task. **(B)** Trial structure for testing trials. **(C)** Levels of stimulus intensity.

### Stimuli

#### Visual Stimuli

As in West et al. (2023), the visual stimuli were Gabor patches. Each Gabor had a Gaussian envelope with a standard deviation of 1.2 degrees of visual angle (dva), a spatial frequency of 0.5 cycles per dva and the starting phase was randomised. The Gabors drifted with a speed of 6 cycles per second.

#### Auditory Stimuli

As in West et al. (2023), the auditory stimuli were pure tones presented in stereo via headphones (Sennheiser HD 202). Each tone included a 5 ms linear onset and offset ramp to avoid clicking artifacts. Tones were synthesised with a 28 kHz sampling rate using an ASIO4ALL sound drive. Loudness was calibrated following the International Standard ISO 226:2003: Acoustics- Normal Equal-Loudness-Level-Contours (International Standardization Organisation, 2003), and final intensity levels were selected through extensive pilot testing.

#### Audio-Visual Stimuli

The audio-visual stimuli were simultaneously presented Gabors and tones, the parameters of which were the same as described above.

### Procedure

All participants completed all combinations of tasks in 3 separate testing sessions (i.e., visual task, auditory task, audio-visual task). The order of the visual and auditory tasks was counterbalanced across participants and all participants completed the audio-visual task in the final session.

Participants were seated in a dark room, at a viewing distance of 57 cm from the screen, and their head was stabilised with a chinrest. Stimuli were presented on a gamma-corrected 60 Hz 1920-by-1080 display. The computer display (ASUS VG248QE Monitor) was connected to a Dell Precision T1700, calibrated with a ColorCal MKII (Cambridge Research Systems). Stimuli were generated and presented using custom code and the Psychophysics Toolbox extensions (Brainard, 1997; Pelli, 1997) for MATLAB.

In each session, participants received instructions for that session’s task which included an explanation about how the stimuli were generated from the relevant category distributions. To further illustrate these distributions, participants were then shown 36 stimuli that were randomly sampled from category 1, followed by category 2. They then completed the category training and testing (detailed below). Each session took approximately 1 hour.

### Testing

At the start of each trial, a fixation cross was presented centrally for 1 s. The fixation cross was then extinguished, and a stimulus was presented (either a visual, auditory or audio-visual stimulus depending on session; all 50 ms in duration). Immediately after the offset of the stimulus, participants were able to respond category 1 or category 2 by pressing a button (F or J keys) with their left or right index finger, respectively. Participants were then prompted to make a confidence judgement for their chosen category with text that read ‘Confidence?’ presented centrally. Participants used the 1-4 number keys to indicate their confidence. The inter-trial interval was 1 s, after which the fixation cross reappeared.

Within a testing block, an equal number of stimuli were shown from each category and intensity level (i.e., 15 trials per cell of the design). Stimulus attributes were sampled randomly from the relevant category distribution. Participants completed 6 testing blocks per session (120 trials per block, a total of 720 trials per session). The order of both category and intensity level was randomised within a block.

At the end of each block, participants were required to take at least a 30 s break. During the break, they were shown the percentage of trials that they had correctly categorised in the most recent block. Participants were also shown a list of the top 10 block scores (across all participants, indicated by initials) for the task they had just completed. This was intended to motivate participants to score highly. We did not provide trial-by-trial feedback during testing to ensure that confidence judgements reflected participants’ subjective evaluations of uncertainty rather than being influenced by external correction signals.

### Training

The trial procedure in training blocks was the same as in testing blocks, except that trial-to-trial feedback was given, only the highest intensity level was used for stimulus presentation and participants were not required to make confidence judgements. For the trial-to-trial feedback, the word “correct” in green or “incorrect” in red was displayed for 1.1 s after participants made their response. Participants completed 2 training blocks (120 trials per block, in total 240 trials) per session. Training was intended to further familiarise participants with the generating category distributions.

## Model Specification

To model responses, we assumed that on each trial, *t*, observers perceived a sensory signal, *x*. Observers transformed that sensory signal into a *decision variable*, the computation of which differed across model classes. To make a discrete category-confidence response, we assumed that observers compared the decision variable to a set of *criteria*. In the following sections, we describe our stimulus standardisation procedure, our assumptions about the sensory signals, decision variable, decision criteria and then describe the model fitting procedure.

### Standardisation of Stimulus Values

To compare stimulus values and model parameters across modalities, we standardised all stimulus values before fitting models. To standardise stimulus values, we subtracted the mean and divided by the standard deviation of all presented stimulus values in each modality such that:

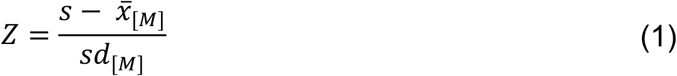

where *Z* is the standardised stimulus value; *s* is the true stimulus value in its original perceptual units (degrees or hertz); 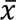 is the mean of all presented stimulus values for a given modality *M* (visual or auditory) and *sd* is the standard deviation of the same set of stimulus values. For the audio-visual task, stimulus values were standardised separately within each modality. This standardisation process was consistent with the idea that participants built up a reliable internal representation of the categories during training such that they understood the relative frequencies of different stimulus values within each category distribution.

### The Sensory Distribution

In all models, we assumed that the observer’s perception of the stimulus, *x*, was a noisy measurement of the true stimulus presented on trial *t* (see **Figure 3A**). We modelled this noisy internal measurement using a Gaussian distribution, referred to as the *sensory distribution*. The sensory distribution was centred on the true stimulus attribute, *s_t_*, with a standard deviation that depended on the intensity of the stimulus, *σ*_*I*_. *σ* was estimated separately at each stimulus intensity level, subject to the constraint that it decreased monotonically as intensity increased. That is, we assumed that greater stimulus intensity was associated with less sensory noise and thus, smaller values of *σ* (see **Figure 3A**). Below we describe the derivation of *x* and σ for both the unidimensional (visual, auditory) and multidimensional (audio-visual) stimuli.

**Figure 3.**
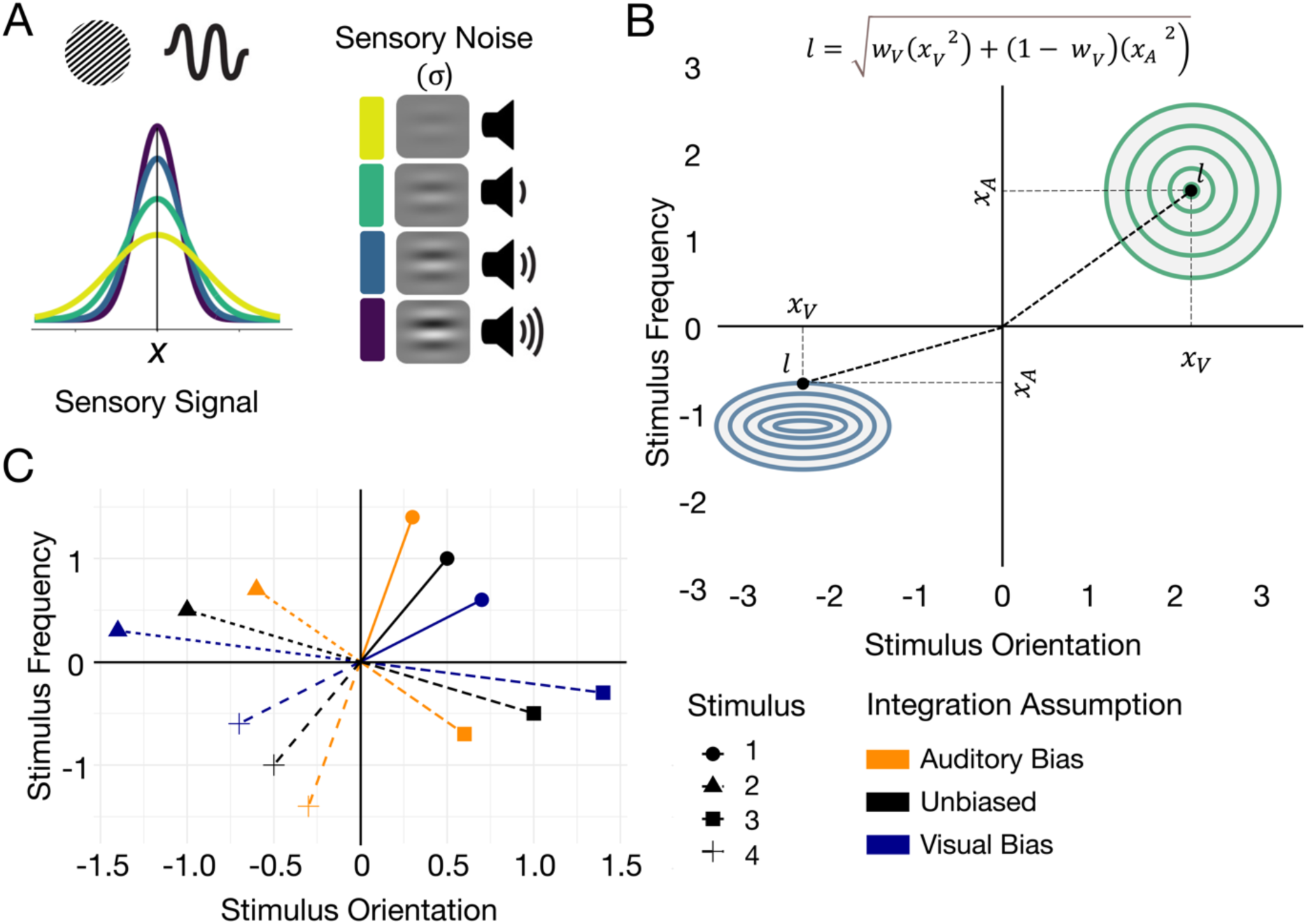
Sensory Signals. *Note.* **(A)** For the unidimensional stimuli, the observers’ internal representation of the stimulus was approximated with a Gaussian distribution centred on the true stimulus attribute with a standard deviation, *σ*, that depended on the intensity level of the stimulus. **(B)** The observer’s internal representation of the multidimensional stimuli was approximated with a bivariate Gaussian distribution. Sensory signals in each dimension, *x*_*A*_ and *x*_*V*_, were combined into a single measure, *l*. Top right quadrant shows unbiased integration of dimensions with the same amount of sensory noise across dimensions. Bottom left quadrant shows unbiased integration with different amounts of sensory noise in each dimension. Samples were taken from the shaded region and each combination of values was transformed into a single measure **(C)** Black data points show unbiased integration of stimulus attributes across dimensions, orange data points show an auditory bias and blue data points show a visual bias for 4 different audio-visual stimuli (shapes).

#### Unidimensional Stimuli

On a given trial, the sensory distribution for the unidimensional stimuli, *x*_*uni*_, was defined as:

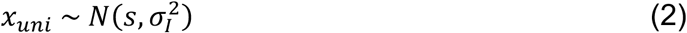

where *N* denotes the normal density function, *s* is the true stimulus attribute, and *σ* is the estimated standard deviation of the distribution, given the stimulus intensity level, *I*.

#### Multidimensional Stimuli

For the multidimensional stimuli, the sensory signal comprised two dimensions, one for each modality. The sensory distribution was thus described by a bivariate Gaussian distribution:

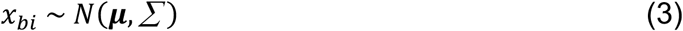

where *N* denotes the normal density function, ***μ*** denotes a vector of stimulus attributes and *∑* a covariance matrix. Specifically, the vector of stimulus attributes on a given trial was described by:

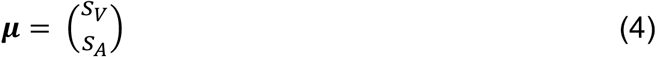

where *s*_*V*_ was the stimulus attribute for the visual dimension and *s*_*A*_ was the stimulus attribute for the auditory dimension. The covariance matrix was described by:

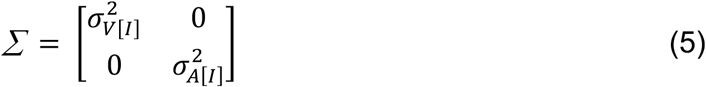

where *σ*_*V*[*I*]_ was the estimated standard deviation of the visual dimension, *σ*_*A*[*I*]_ was the estimated standard deviation of the auditory dimension, given the stimulus intensity level, *I*. We tested two different assumptions about *σ*: either that *σ* differed across the stimulus dimensions, referred to as the *different noise* variant, or that *σ* was the same across the stimulus dimensions, referred to as the *same noise* variant.

##### Same Noise

For the *same noise* model variants, we assumed that sensory noise was the same across the two stimulus dimensions such that:

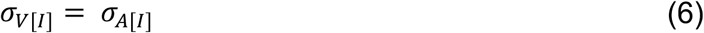

where *σ*_*V*_ was the estimated amount of sensory noise in the visual dimension, *σ*_*A*_ was the estimated amount of sensory noise in the auditory dimension, and *I* indexed the stimulus intensity level (see top right panel of **Figure 3B**).

##### Different Noise

For the *different noise* variants, we assumed that sensory noise differed across the two stimulus dimensions such that *σ*_*V*[*I*]_ ≠ *σ*_*A*[*I*]_, where *σ*_*V*_ was the estimated amount of sensory noise in the visual dimension, *σ*_*A*_ was the estimated amount of sensory noise in the auditory dimension and *I* indexed the stimulus intensity level (see bottom left panel of **Figure 3B**). Under the different noise assumption, we estimated *σ* separately for each stimulus dimension and intensity level, constrained by monotonicity within each domain.

##### Transformation of Bivariate Sensory Distribution

For ease of exposition, we transformed the bivariate sensory distribution, *x*_*bi*_, into a univariate Gaussian distribution defined by:

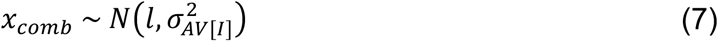

where *N* denotes the normal density function, *l* denotes the mean and *σ*_*AV*_ the standard deviation of the resulting univariate distribution. We calculated *l* as the Euclidian distance of a set of multidimensional stimulus coordinates, [*x*_*V*_, *x*_*A*_], from the centre of the standardised stimulus space, [0,0] (henceforth referred to as the *origin*). We tested two different assumptions about the computation of *l*: either that the observer attended to both dimensions of the stimulus (*integration* model variants) or that the observer attended to only a single dimension of the stimulus (*max* model variants).

##### Integration Variants

For the integration model variants, we assumed that the observer attended to both dimensions of the stimulus to make their decision. We therefore computed a weighted combination of the sensory signals received from each dimension such that:

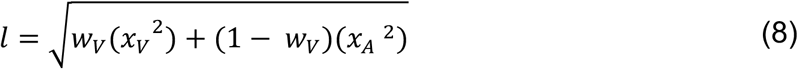

where *w*_*V*_ was the weight for the visual stimulus dimension, *x*_*V*_ was the perceived stimulus value for the visual dimension and *x*_*A*_ was the perceived stimulus value for the auditory dimension. See **Figure 3C** where orange data points show integration with an auditory bias (where *w*_*V*_ < 0.5) and blue data points show integration with a visual bias (where *w*_*V*_> 0.5). The weights were constrained to be non-negative and bounded on the interval [0, 1] such that *w*_*V*_ + *w*_*A*_ = 1.

##### Max Variants

For the max model variants, we assumed that the observer attended to a single dimension of the stimulus only and thus, the computation of *l* depended on a single value from the set of stimulus coordinates [*x*_*V*_, *x*_*A*_]. The “selected” dimension was determined by which value was the most informative about the generating category. The informativeness of the generating category was referred to as *category diagnosticity* and reflected the probability density of the presented stimulus value for the most probable category distribution, normalised by the total probability density of the stimulus for both category distributions:

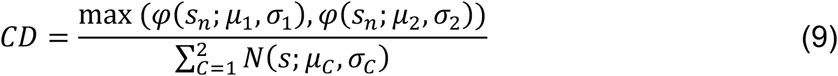

where *φ* denoted the normal density function, *μ*_1_ and *μ*_2_ were the means of the generating category 1 and category 2 distributions and *σ*_1_ and *σ*_2_ were the standard deviations of the generating category 1 and category 2 distributions. *C* denoted the category and *s*_*t*_ was the stimulus attribute on trial *t*. The distance, *l*, was then calculated as the absolute value of the selected dimension:

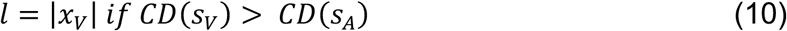

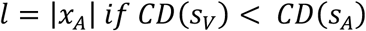

The computation of *l* for the max models was thus consistent with the integration models because the result was equivalent to a distance measure involving only a single term.

###### Same Noise and Different Noise Variants

The standard deviation of *x_comb_*, *σ*_*AV*_, depended on the model variant, specifically the same noise or different noise assumption *and* the integration or max assumption. Under the same noise assumption:

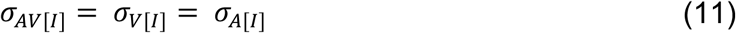

where *σ*_*V*_ was the estimated standard deviation of the visual dimension of *x_bi_*, *σ*_*A*_ was the estimated standard deviation of the auditory dimension of *x_bi_*, and *I* indexed the stimulus intensity level.

Under the *integration different noise* assumption, we calculated a weighted combination of the standard deviations of each dimension of the bivariate sensory distribution *x_bi_*:

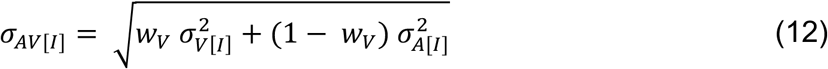

where *w*_*V*_ was the estimated weight for the visual dimension of the stimulus (as in Equation 8), *σ*_*V*_ was the estimated standard deviation of the visual dimension of *x_bi_*, *σ*_*A*_ was the estimated standard deviation of the auditory dimension of *x_bi_*, and *I* indexed the stimulus intensity level. Using Equation 12 to combine the estimates of *σ* across modalities meant that the resulting value of *σ*_*AV*_ was of the same magnitude as *σ*_*V*_ and *σ*_*A*_. Under the *max different noise* assumption,

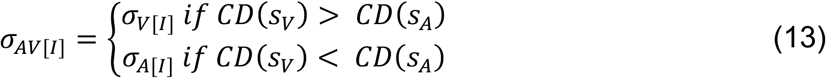

where *CD*(*s*_*V*_) was the category diagnosticity of the visual dimension of the stimulus and *CD*(*s*_*A*_) was the category diagnosticity of the auditory dimension of the stimulus (see Equation 9).

### Decision Variable

In all models, we assumed that the observer transformed the sensory signal into a *decision variable* which was then compared with a set of criteria. We tested different assumptions about the computation of the decision variable which quantified different theoretical models about the choice and confidence generation process. We categorised these theoretical models into three broad classes: (1) the unscaled evidence strength models, (2) the scaled evidence strength models and (3) the Bayesian models. One of the distinguishing factors among candidate classes concerned whether the decision variable was represented in *psychological space* (unscaled evidence strength and scaled evidence strength; discussed in more detail below) or *probability space* (Bayesian; discussed in more detail below).

For models that represented the decision variable in *psychological space*, the decision variable was assumed to be in the same psychological units as the sensory signals. For these models, we therefore use *x* to denote both the sensory signal and the decision variable. For models that represented the decision variable in *probability space*, we assumed that the sensory signal was transformed into a probability metric which was then compared with choice and confidence criteria in the same probability units. For these models we use *d* to denote the decision variable and distinguish it from the sensory signal.

To illustrate the difference between the computation of a decision variable which is represented in psychological space or probability space: for a non-Bayesian model with category-confidence criteria in psychological space, a Category 2 Confidence 1 response might be made for any visual stimulus perceived as having an orientation between 0 and 5 degrees (or 0 and 0.6 in standardised stimulus units). For a Bayesian model, where the criteria were in probability space, the same Category 2 Confidence 1 response might be made for any visual stimulus perceived as having a log posterior probability ratio between 0 and 0.5.

#### Non-Bayesian Models

As described above, for the non-Bayesian models, the sensory distribution and the decision variable distribution were represented in the same psychological space. This meant that *the sensory distributions* (*x_uni_* and *x_comb_* for the unidimensional and multidimensional stimuli respectively) *were equivalent to the decision variable distributions*.

#### Bayesian Models

Consistent with Bayesian decision theory, for the Bayesian models we assumed that the observer combined prior knowledge of the generating category distributions with observed stimulus information to form posterior beliefs about category membership. For all Bayesian models, therefore, we assumed that the observer transformed the sensory signal into a log posterior probability ratio, *d*, to make a decision. The log posterior probability ratio represented the (log) ratio of posterior beliefs about category membership, such that 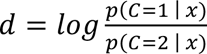 where *x* represented the perceived value of the stimulus on a given trial. When the log posterior probability ratio was positive, there was greater evidence for category 1. For the Bayesian models, we assumed that category and confidence reports depended on the sensory signal *x* only via *d.* Following Adler and Ma (2018), we used the following derivation of *d:*

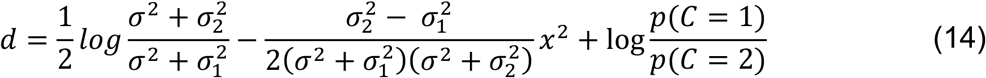

where *σ*_1_ was the standard deviation of the category 1 distribution, *σ*_2_ was the standard deviation of the category 2 distribution and σ was the fitted value of sensory noise, approximating the amount of sensory uncertainty in the observer’s internal representation of the stimulus. Because we sampled equally from each category, we assumed that *p*(*C*=1)=*p*(*C*=2)=0.5 such that 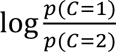 was 0 in Equation 14.

##### Log Posterior Probability Ratio with Free Category Distribution Parameters

The log posterior probability ratio model described above assumes that the observer knows the true values of the standard deviations of the category distributions *σ*_1_ and *σ*_2_ in Equation 14. We also tested a version of the model in which the observer had imperfect knowledge of the parameters of the generating category distributions. In this model, we estimated the values of *σ*_1_ and *σ*_2_ as free parameters.

### Choice and Confidence Criteria

As described above, in all models we assumed that the decision variable was compared with a set of criteria. The criteria partitioned the range of the decision variable into discrete category and confidence response regions so that if the decision variable fell within a defined region, response *r* was given (see **Figure 4**). Given four confidence levels for each of the two categories, this yielded eight response regions, *r* ∈ {1, 2, 3, …, 8}, each corresponding to a unique category and confidence combination. The choice and confidence criteria were positioned in the same units as the decision variable and required different assumptions across the different theoretical model classes. These assumptions are described below for the non-Bayesian models, where the decision variable was measured in psychological space, and Bayesian models, where the decision variable was measured in probability space.

**Figure 4.**
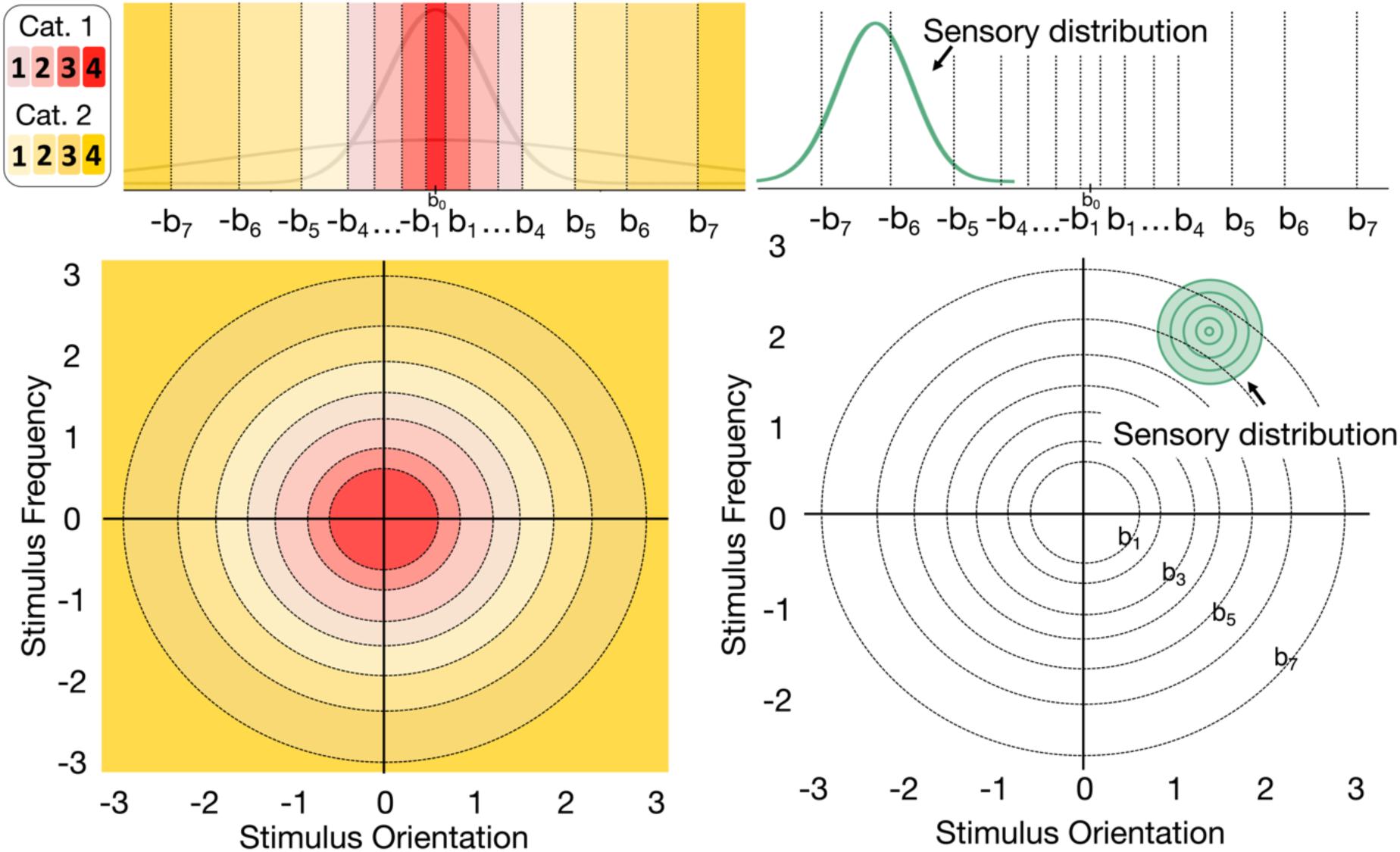
Choice and Confidence Criteria Across Unidimensional and Multidimensional Tasks. *Note.* In the unidimensional task (top panel), 7 symmetrical criteria partitioned the range of the decision variable into discrete category and confidence response regions (top left panel; see legend at the bottom for corresponding responses). The sensory distribution was compared against the choice and confidence criteria to calculate a response probability (top right panel). In the multidimensional task, 7 circular criteria partitioned the response regions (bottom left panel) and were compared against the bivariate sensory distribution (bottom right panel).

#### Non-Bayesian Choice and Confidence Criteria

The non-Bayesian models constituted models from both the scaled evidence strength and unscaled evidence strength class. For all these models, we refer to the position of the criteria in psychological space as *b*. To define the eight response regions across the range of psychological space, we used seven criteria. We assumed that each criterion increased monotonically across successive criteria such that higher confidence response regions were aligned with more extreme category evidence (see **Figure 4**). We also assumed that the criteria were positioned symmetrically around the means of the category distributions. We refer to this centre point as *b*_0_. We assumed the criteria were symmetrical in this task because two measurements that are equidistant from the middle of the category structure but on opposite sides of *b*_0_ should logically have the same confidence associated with them. Due to symmetry, confidence regions on equidistant but opposite sides of *b*_0_ were treated as equivalent by the model. The criteria were ordered such that:

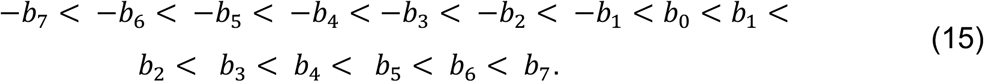

The symmetrical criteria −*b*_4_ and *b*_4_ divided the stimulus range into two different category response regions such that:

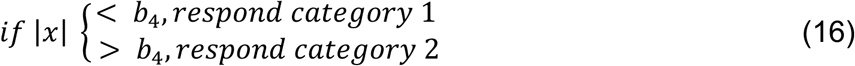

To make a confidence response, we assumed that the observer used 4 symmetrical criteria for each category: *b*_1_– *b*_4_ for category 1 responses and *b*_4_ - *b*_7_ for category 2 responses. See **Figure 4A** to see how the criteria aligned with category evidence. Thus, the observer used a decision rule such that:

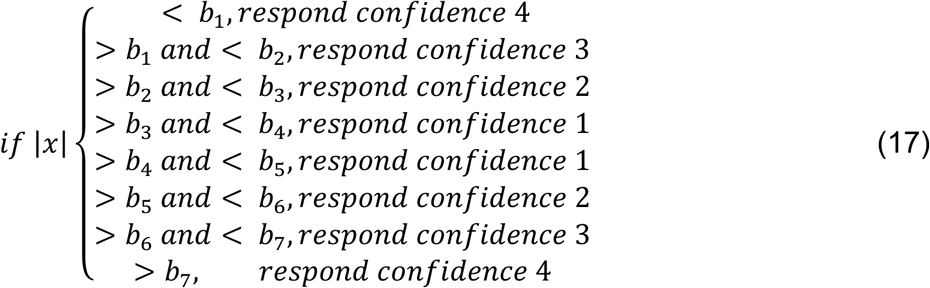

For the unscaled evidence strength models and the scaled evidence strength models, we made different assumptions about how *b* varies as a function of sensory uncertainty.

##### Unscaled Evidence Strength Model

In the unscaled evidence strength model, the choice and confidence criteria were not dependent on (estimated) values of sensory uncertainty (*σ*) and therefore occupied fixed positions across intensity levels. The position of each criterion in psychological space was described by a free parameter, *k*. The resulting criteria in psychological space were given by:

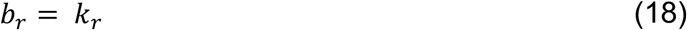

##### Scaled Evidence Strength Model

In the scaled evidence strength model, the positions of the choice and confidence criteria were functions of *σ*. Each criterion depended on 4 free parameters: *k*, *m, z* and *σ*. *k* represented the base position of each criterion which was offset by *mσ*^*z*^, where *m* was also estimated separately for each criterion. The resulting criteria in psychological space were given by:

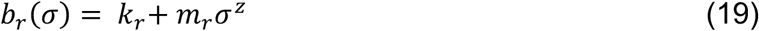

We constrained *z* to be greater than 1. Previous research has also considered different scaling rules (Adler & Ma, 2018; Locke et al., 2022; West et al., 2023), for example where criteria are estimated as linear or quadratic functions of *σ*. We chose not to investigate these variants directly here to reduce the number of models for model comparison and importantly, our chosen evidence strength model contained both linear and quadratic scaling as special cases.

#### Bayesian Choice and Confidence Criteria

For the Bayesian models, we assumed that the observer chose a response by comparing *d,* the log posterior probability ratio for a given stimulus (see Equation 14), with a set of choice and confidence criteria *k* = (*k*_1_, *k*_2_ … *k*_7_) that partitioned *d* space into eight response regions, each representing a unique category and confidence combination. The positions of the choice and confidence criteria for the Bayesian models, therefore, were free parameters in *d* units. We assumed that each criterion increased monotonically across successive criteria such that higher confidence response regions aligned with greater absolute values of *d* (**Figure 4D**). Thus, the choice and confidence criteria were ordered such that:

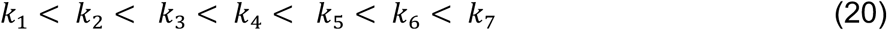

*k*_4_ divided the stimulus range into two different category response regions such that

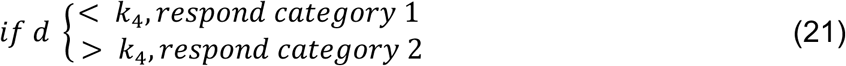

For confidence response, the observer used a decision rule such that:

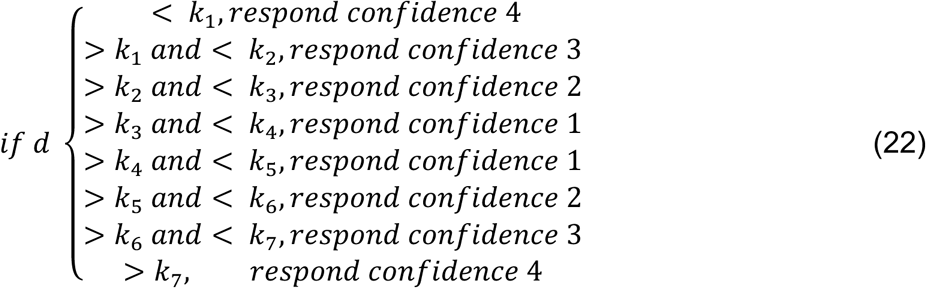

### Fitting the Models to Data

Model parameters were estimated for each participant using maximum likelihood estimation. Below we first describe the process for estimating the parameters of the bivariate sensory distribution and then describe the process for calculating the likelihood of a given response, *r*, for a model with parameters *θ*. We describe this process separately for the non-Bayesian and Bayesian models, as they required different transformations.

#### Bivariate Sensory Distribution

To estimate the parameters of the bivariate sensory distribution, we sampled 10 evenly spaced values from the 5^th^ to 95^th^ percentile of the cumulative density distribution for each dimension of the sensory distribution. We generated all possible combinations of these sampled values (see shaded region of bottom left panel of **Figure 3B**) and transformed each combination into the relevant univariate decision variable, *x*_*comb*_, given the model variant assumptions (integration vs. max and different noise vs. same noise). See above for detailed descriptions for each set of assumptions. For each univariate Gaussian decision variable, we computed a likelihood (described below) which was then multiplied by the normalised joint density of the sample, given the bivariate sensory distribution, *x*_*bi*_.

#### Likelihood Calculation: Non-Bayesian Models

For the non-Bayesian models, the decision variable distributions, *x*_*uni*_ and *x*_*comb*_, were in the same psychological units as the choice and confidence criteria, *b*, and thus, could be compared directly. For each trial, we therefore determined the predicted response probability for a chosen combination of category and confidence level by computing the probability mass of the sensory distribution between adjacent criteria, given the participant’s response on that trial, *r*_*t*_(cf. Luce, 1959; see Fig 3D). This quantity was defined as:

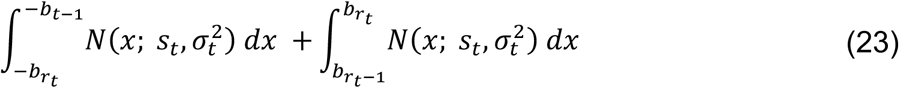

Where *b*_0_ = 0, *b*_8_ = ∞ and − *b*_8_ = −∞.

#### Likelihood Calculation: Bayesian Models

In the Bayesian models, the choice and confidence criteria were positioned in probability space (log posterior probability ratio units) and thus, the sensory distribution and criteria could not be compared directly. We, therefore, transformed the choice and confidence criteria into psychological space by using the relevant value of *d* as the left-hand side of Equation 14 and solved for *x*. Solving Equation 14 for *x* requires calculating the square root of *d*, which is undefined for negative values. We therefore used the absolute value of *d* to solve Equation 14 for *x* and applied the appropriate sign to the result to arrive at a final standardised value for *x*. Where the category distribution parameters were estimated as free parameters, we used the estimated values in place of the true standard deviations (*σ*_1_ and *σ*_2_). The resulting choice and confidence criteria in psychological space were then compared with the decision variable distribution, described below.

##### Unidimensional Stimuli

After transforming the choice and confidence criteria in *d* space to psychological space using Equation 14, we computed the probability mass of the sensory distribution between adjacent criteria using Equation 23.

##### Multidimensional Stimuli

In order to transform the choice and confidence criteria into psychological space we used the value of *d* as the left-hand side of Equation 14 to solve for *x*. For the integration variants, we did this separately for each stimulus dimension such that we used the estimated value of *σ* and the standard deviations of the generating category distribution (*σ*_1_ and *σ*_2_) for each dimension. This gave us a “threshold” value in psychological space for each stimulus dimension, referred to as *y_V_* and *y_A_*. These values were then converted into a combined distance metric, *b_comb_*, according to:

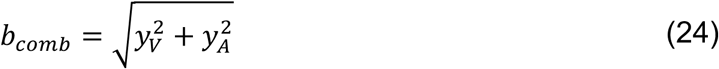

*b_comb_*, therefore, represented a criterion in psychological space calculated as a combination of the “threshold” values for each stimulus dimension. The criteria were unweighted, as the weighting was applied to the stimulus values directly (see Equation 8). For the max variants, we converted the estimated choice and confidence criteria into psychological space, see Equation 14, for the selected stimulus dimension only, see Equation 10.

#### Log Likelihood of the Dataset

To obtain the log likelihood of the dataset, we computed the sum of the log probability for every trial *t,* where *T* is the total number of trials.

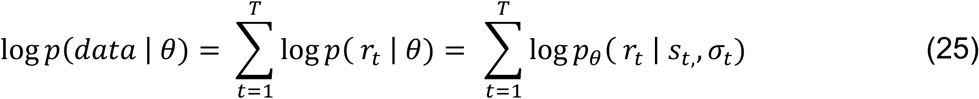

### Model Selection

To compare candidate models, we used Akaike information criterion (*AIC*) and Bayesian information criterion (*BIC*) for model selection. For a single participant, *AIC* is defined as:

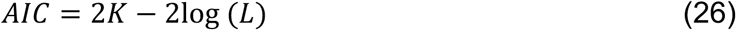

where *K* is the number of model parameters and log(*L*) is the log likelihood of the dataset. To evaluate the performance of each model across all participants, we summed the *AIC* values across the *N* participants,

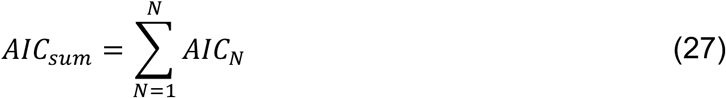

To compare the relative support for each model, we also calculated Akaike weights (Wagenmakers & Farrell, 2004). First, for each model *M*_*m*_, where *m* = 1, 2, …, *J*, we computed the difference in *AIC* relative to the best candidate model (the model with the lowest *AIC* value):

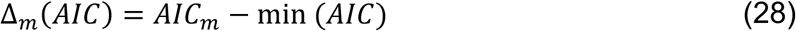

Next, we calculated the Akaike weight for each model as:

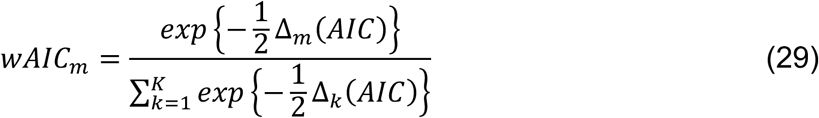

The Akaike weights can be interpreted as the probability that a particular model is the best model in the set of candidate models, given the data and specified models (Wagenmakers & Farrell, 2004).

For a single participant, *BIC* is defined as:

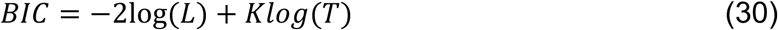

where *K* is the number of model parameters, log(*L*) is the log likelihood of the dataset and *T* is the number of trials in the dataset. Because *BIC* is sensitive to the total sample size used to compute the likelihood, calculating the summed *BIC* for each model requires that the sample size accounts for the total number of trials across all participants. We therefore summed the likelihood terms across participants, *N*, and added to that the total number of free parameters across participants, *kN*, multiplied by the logarithm of the total number of trials across participants, *tN*:

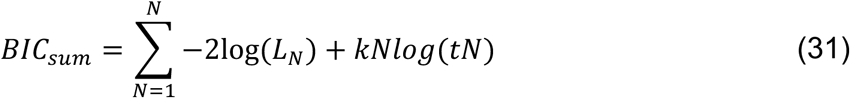

### Parameter and Model Recovery

We performed both parameter and model recovery for our core models. In the case of parameter recovery, we simulated data using known parameter values, fit the models to the simulated data, and then checked the correlation between the true generating parameters and the parameters estimated during the fitting procedure. For model recovery, we simulated datasets from each model and fit all models to these datasets. We then evaluated the probability that the model used to generate the data was correctly identified as the best-fitting model based on a specific model selection metric (*AIC* or *BIC*). Model recovery was conducted at two levels: individual model variants (see **Figure 5A**) and broader model classes (see **Figure 5B**). Evaluating model recovery at the model class level allowed us to determine how well the core theoretical assumptions distinguishing different groups of models aligned with the data (West et al., 2023). For both the parameter and model recovery, we used parameter values that were similar to the range of values obtained through fitting participant data. Further, to match our empirical dataset, we performed the parameter and model recovery with simulated datasets consisting of 10 ‘participants’ and 720 trials per participant.

**Figure 5.**
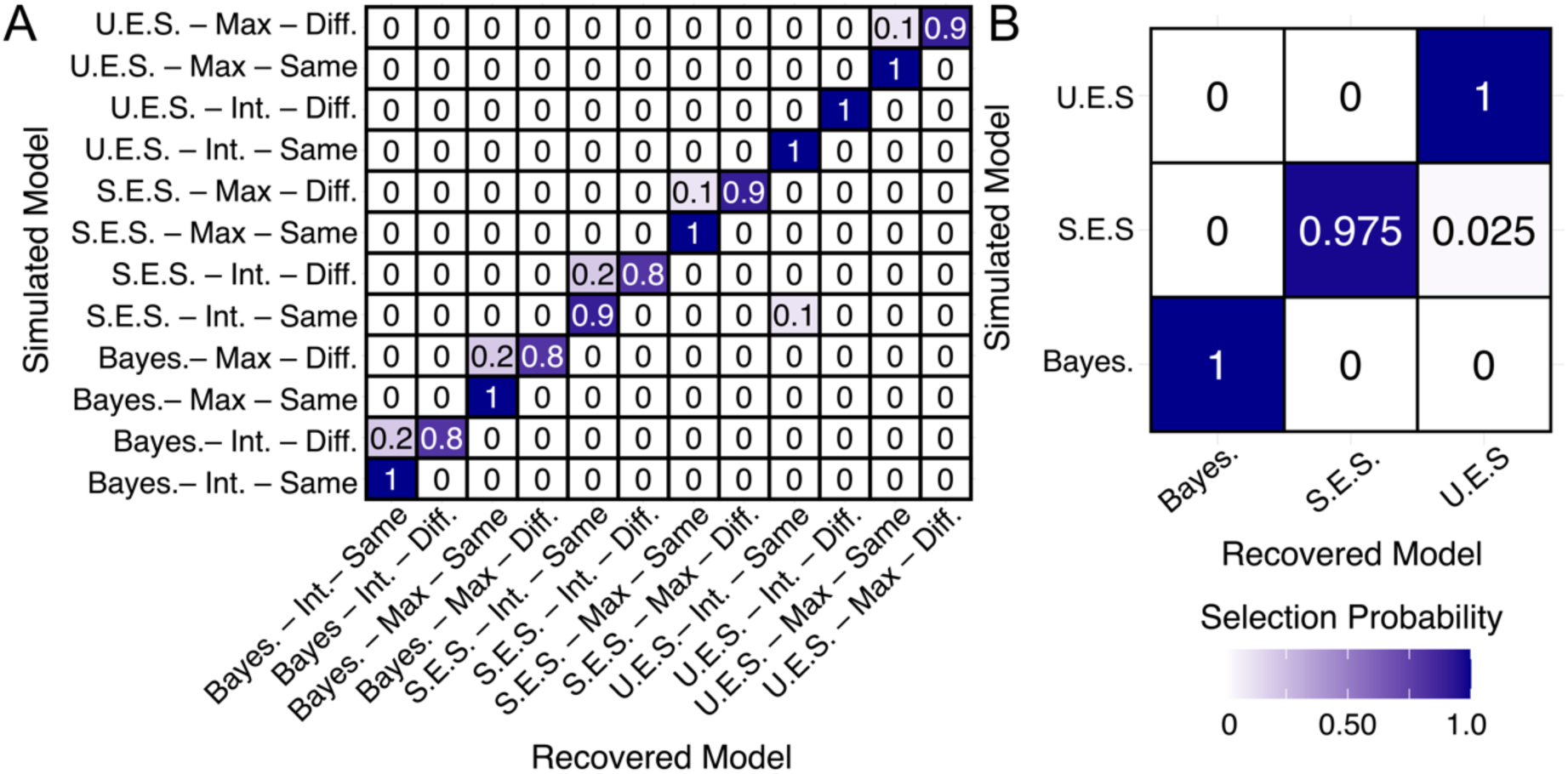
Model Recovery. *Note.* Numbers and colours denote the probability that the data generated with model X (y-axis) are best fit by model Y (x-axis). Confusability matrix for **(A)** all model variants and **(B)** model classes using *AIC* for model selection. U.E.S refers to the unscaled evidence strength model class. S.E.S refers to the scaled evidence strength model class. Bayes. refers to the Bayesian model class. Int. refers to the integration assumption. Diff. and same refers to the assumption about the noise parameters across modalities.

As shown in **Table S 3 – Table S 5** and **Figure S 3 – Figure S 5**, parameters generally exhibited strong recovery across models, with the vast majority of correlation coefficients exceeding *r* > 0.75 (White et al., 2018). However, there were some exceptions: noise parameters (*σ*) in the unscaled evidence strength and Bayesian models showed poorer recovery (*r* < 0.5) under different noise and integration assumptions. The poorer recovery could have stemmed from using a restricted range of parameter values when simulating data. For the unscaled evidence strength and Bayesian models in particular, small changes in the noise parameters often resulted in minimal changes to the model’s predictions of category and confidence responses, making parameter recovery in these cases more difficult. This issue would have likely been more pronounced as the number of parameters in the model increased (as in the different noise and integration assumptions).

As shown in **Figure 5B**, model class recovery was robust overall. This was particularly true when using *AIC* for model selection (see **Figure S 6B),** which aligns with findings from our previous work (West et al., 2023). For the bidimensional models, recovery was also strong across assumptions (see **Figure 5A**), though the integration models with the same noise assumption were slightly over-selected. This is likely because we simulated with similar noise values across modalities, so the additional four parameters in the different noise models did not improve the model’s fit sufficiently to outweigh the penalty for added complexity. Using *AIC* for model selection also led to better recovery for the bidimensional models (see **Figure S 6A**).

## Results

### Participant Exclusion

Following the approach in West et al. (2023), we excluded one participant whose categorisation accuracy was not significantly above chance (50%) in the highest intensity condition, as determined by a binomial test of significance (see **Table S 1**).

### Confirming the Effect of Stimulus Manipulations: Model Free GLMMs

Following the approach in West et al. (2023), we first verified that our stimulus manipulations had the desired effect on category and confidence responses across tasks. To do so, we used generalised linear mixed models (*GLMM*s) to predict category and confidence responses from the relevant stimulus features for each task. Since the expected relationships between stimulus attributes and responses were non-linear, we first transformed the raw stimulus attributes. For category responses, this involved calculating a measure of ‘evidence for category 2’.

#### Confirming that the stimulus manipulations affected category responses

For each task, we calculated evidence for category 2 by determining the probability density of the presented stimulus attribute under the category 2 distribution, normalised by the combined probability density across both category distributions:

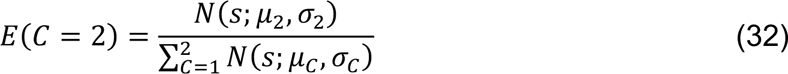

where s is the value of the stimulus, *C* is the category and *N* denotes the normal density function. The result indicated ‘evidence for category 2’ for each stimulus attribute, which could range from 0.2 (stimulus was highly likely to be from category 1) to 1 (stimulus could be from category 2 only). The evidence value never reaches 0 because the category 1 distribution falls entirely within the range of the category 2 distribution (see **Figure 2A**).

We fit linear mixed models using a logistic link function to predict category response (1 or 2) from stimulus intensity and evidence for category 2, for each task and modality. The models included an intercept term, intensity, evidence and the interaction term between intensity and evidence as random effects. For each regression analysis, we report the odds ratio for the fixed effect and 95% confidence intervals in **Table 1**. This analysis was conducted to descriptively assess the effects of stimulus manipulations on responses across tasks so additional statistical details and follow-ups are provided in **Table S 2**.

**Table 1.**
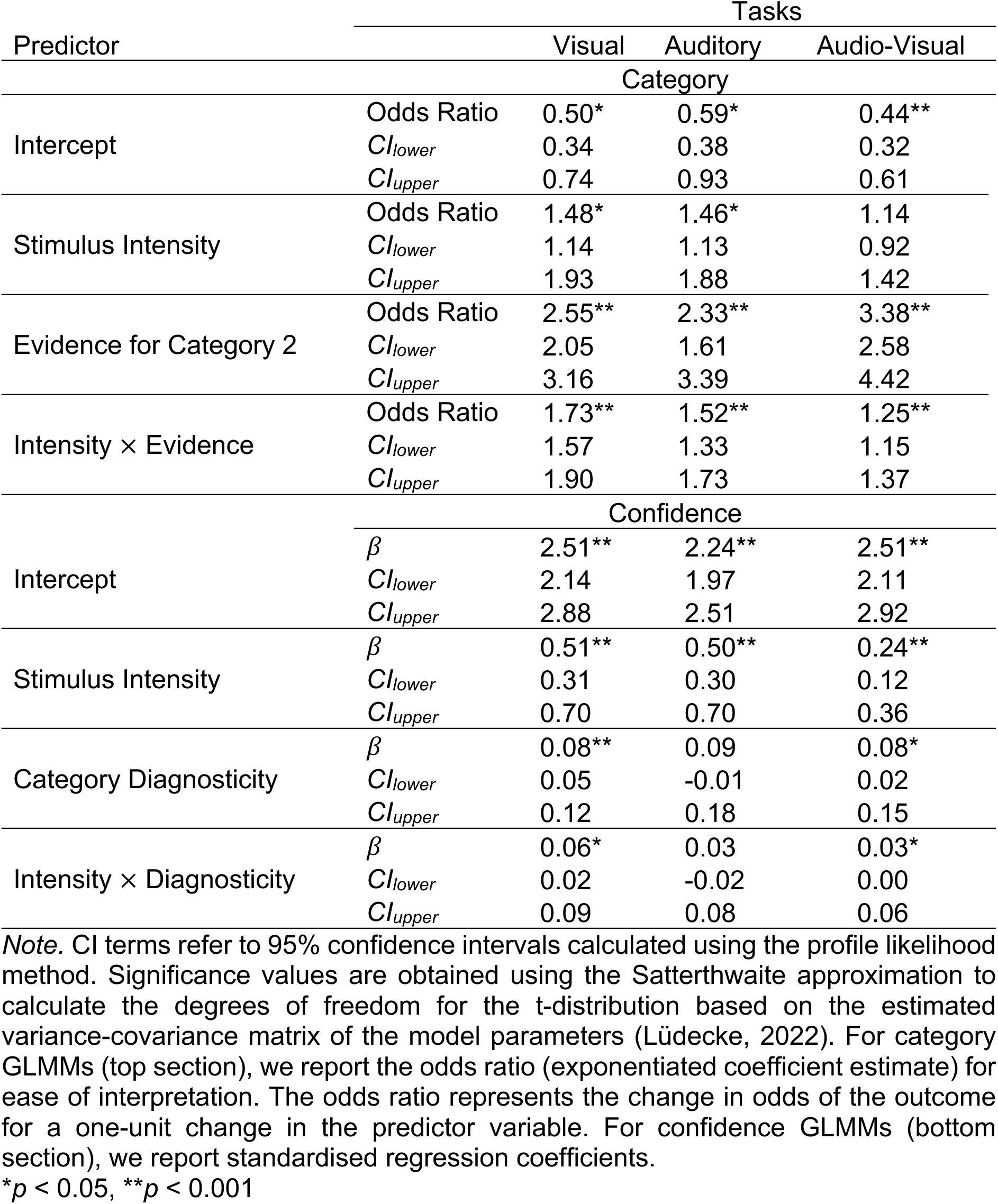
Model-Free GLMMs for Category and Confidence Responses.

As expected, stimulus values that indicated increasing evidence for category 2 were positively predictive of category 2 responses in the visual (*OR_visual_* = 2.55, 95% CI [2.05-3.16], *p* < .001), auditory (*OR_auditory_* = 2.33, 95% CI [1.61-3.39], *p* = < .001), and audio-visual tasks (*OR_audio-visual_* = 3.38, 95% CI [2.58-4.42], *p* < .001). Increasing stimulus intensity was also positively predictive of category 2 responses for the visual (*OR_visual_* = 1.48, 95% CI [1.14-1.93], *p* = .004) and auditory tasks (*OR_auditory_* = 1.46 [1.13-1.88], *p* = .004) but not the audio-visual task (*OR_audio-visual_* = 1.14, 95% CI [0.92-1.42], *p* = .223). These main effects were qualified by significant two-way interactions between category 2 evidence and intensity, for the visual (*OR_visual_* = 1.73, 95% CI [1.57-1.90], *p* = < .001), auditory (*OR_auditory_* = 1.52, 95% CI [1.33-1.73], *p* < .001) and audio-visual tasks (*OR_audio-visual_* = 1.25, 95% CI [1.15-1.37], *p* < .001). As illustrated in **Figure 6A**, the effect of increasing category 2 evidence on category 2 responses was strongest for the highest intensity stimuli and decreased with stimulus intensity (see **Table S 2** for follow up tests).

**Figure 6.**
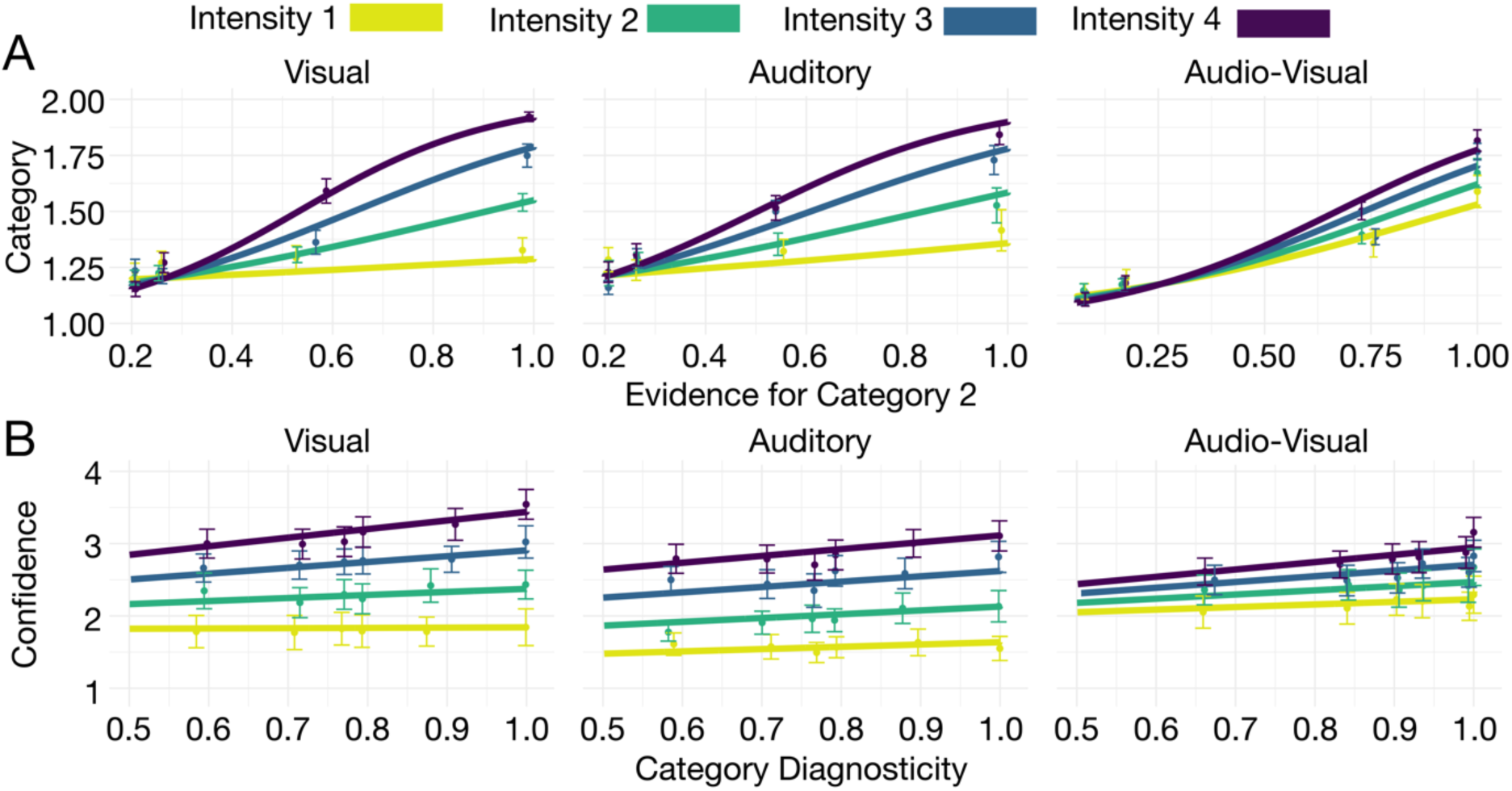
Model Free GLMMs. *Note***. (A)** Predictions from a *GLMM* on category responses showed that the proportion of category 2 responses increased as a function of evidence for category 2 across the visual, auditory and audio-visual tasks (columns). The strength of this relationship was modulated by stimulus intensity (colours). **(B)** Predictions from a *GLMM* on confidence responses showed that across all tasks (columns), confidence increased as category diagnosticity, the relative evidence for the most probable category, increased. The strength of this relationship was modulated by stimulus intensity (colours).

These findings confirmed that participants understood the category distributions across modalities. Specifically, as the relative probability of a stimulus being sampled from category 2 increased, participants were more likely to categorise that stimulus as being sampled from category 2. The influence of category 2 evidence was modulated by stimulus intensity. At lower intensities, responses were more random and less guided by the category distributions. This pattern was consistent across tasks, confirming that the stimulus manipulations had the intended effect on category responses.

#### Confirming that the Stimulus Manipulations Affected Confidence

To compare the effect of the stimulus manipulations on confidence across the different tasks we transformed stimulus values to ‘category diagnosticity’ using Equation 9 prior to fitting the GLMMs. The result indicated category diagnosticity for each stimulus value, which could range from 0.5 (both categories are equally likely) to 1 (stimulus could be from the most probable category only). We fit linear mixed models to predict confidence from stimulus intensity and category diagnosticity for each modality. The models included an intercept term, intensity, diagnosticity and the interaction term between intensity and diagnosticity as random effects. For each regression analysis, we report the regression coefficient for the fixed effect and 95% confidence intervals in **Table 1**.

Increasing category diagnosticity was predictive of increasing confidence in the visual (*β*_*visual*_= 0.08, 95% CI [0.05-0.12], *p* < .001) and audio-visual tasks (*β*_*audio*–*visual*_= 0.08, 95% CI [0.02-0.15], *p* = 0.10), but not the auditory task (*β*_*auditory*_ = 0.09, 95% CI [-0.01-0.18], *p* = .066). Increasing stimulus intensity was predictive of increasing confidence in the visual (*β*_*visual*_= 0.51, 95% CI [0.31-0.70], *p* < .001), auditory (*β*_*auditory*_ = 0.50, 95% CI [0.30-0.70], *p* < .001) and audio-visual tasks (*β*_*audio*–*visual*_ = 0.24, 95% CI [0.12-0.36], *p* < .001). We also observed significant two-way interactions between category diagnosticity and intensity in the visual (*β*_*visual*_= 0.06, 95% CI [0.02-0.09], *p* = .002) and audio-visual task (*β*_*audio*–*visual*_= 0.03, 95% CI [0.00-0.06], *p* = 0.024), but not the auditory task (*β*_*auditory*_= 0.03, 95% CI [-0.02-0.08], *p* = .211). For the visual and audio-visual task, increasing category diagnosticity was associated with increasing confidence but the strength of this effect was modulated by stimulus intensity. As illustrated in **Figure 6B**, the effect of increasing category diagnosticity was strongest for the highest intensity stimuli and decreased with lower stimulus intensities (see **Table S 2** for follow up tests).

These findings indicate that as the relative evidence for a given category increased, participants’ confidence in their chosen category increased, especially for the visual and audio-visual tasks. When combined with the category GLMMs, which showed that participants’ category responses corresponded with the most probable category, these results indicate that participants understood the category distributions and used them to inform their confidence responses. The effect of the category distributions on confidence was modulated by stimulus intensity, with lower stimulus intensity leading to uniformly low confidence responses. In the auditory task, changes in confidence were driven primarily by changes in stimulus intensity and not category diagnosticity.

### Computational Modelling

The GLMMs confirmed that our chosen stimulus manipulations had the expected effects on category and confidence responses across modalities. To further explore the underlying computations driving these judgements, we used computational modelling. We assessed the performance of confidence models from three classes: 1) unscaled evidence strength models, 2) scaled evidence strength models and 3) Bayesian models. To identify shared underlying algorithms, we first tested whether a single class of models provided the best fit across modalities. In the following sections, we briefly revisit the assumptions of each model class and describe their performance across modalities.

### Performance of Model Classes

#### Unscaled Evidence Strength Model

According to the unscaled evidence strength account, confidence reports depend on the distance between the sensory measurement and the category decision criterion (*evidence strength*) only. These models do not independently quantify *sensory uncertainty* as a core part of the decision process. Specifically, for the unscaled evidence strength model, we assumed that the observer compares their internal representation of a stimulus to a set of choice and confidence criteria that do not change with stimulus intensity.

As a result of the fixed choice and confidence criteria, the model predictions remain unchanged across intensity levels, as shown in **Figure 7A**. On the contrary, the empirical confidence ratings clearly change as a function of stimulus intensity, leading to a poor fit of the model to data across tasks. In line with this qualitatively poor fit, the unscaled evidence strength model performed worst in terms of summed *AIC* and *BIC* (see **Figure 7A**) and was selected as the best fitting class for only a single participant in the auditory and audio-visual task and for no participants in the visual task (see **Figure 9A**). These findings suggests that a model in which confidence depends only on the strength of the evidence, and not on the amount of sensory uncertainty in the representation of the stimulus, fails to accurately account for the cognitive processes involved in perceptual metacognition.

**Figure 7.**
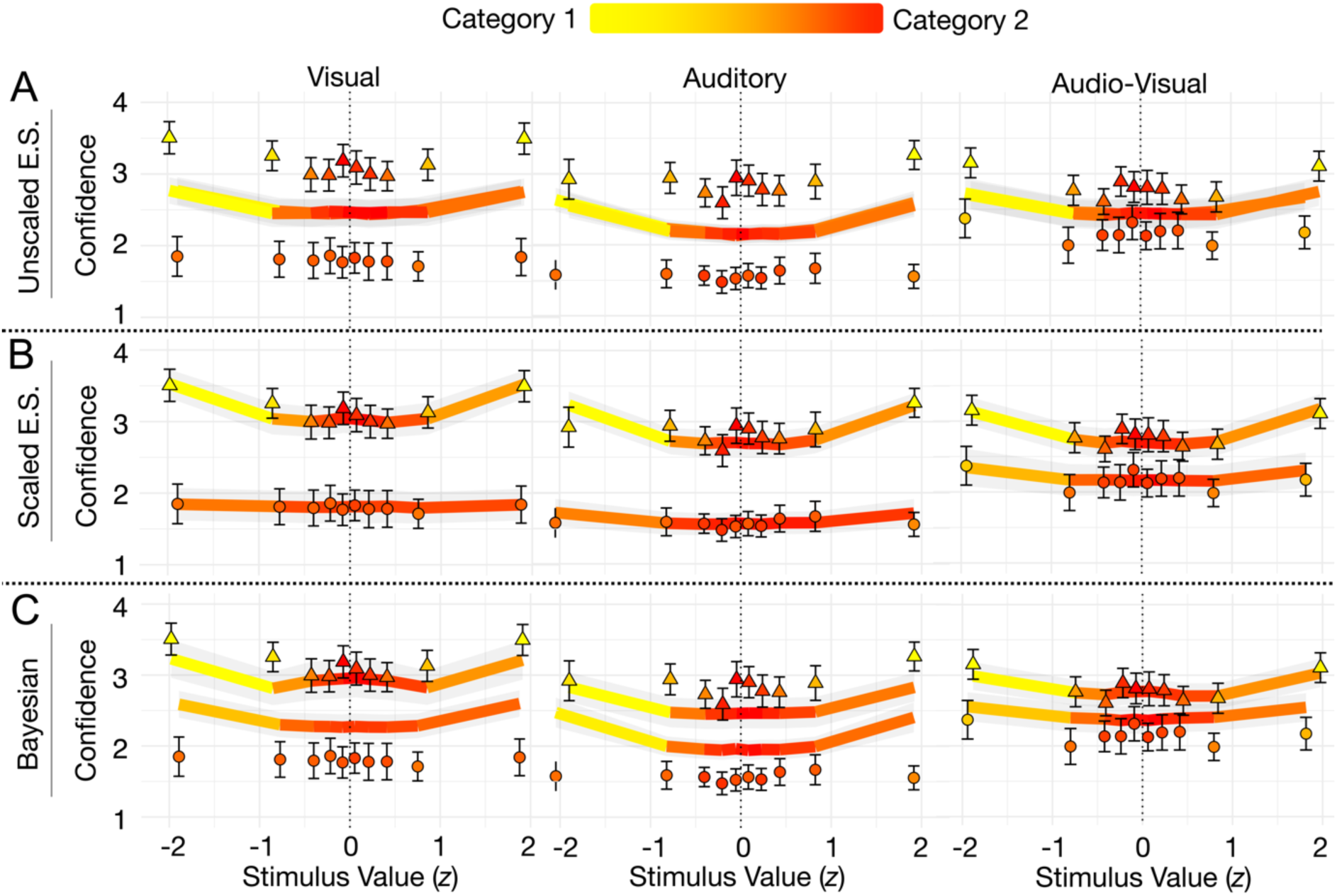
Model Predictions Across Classes for Each Task. *Note.* Columns show task modalities. Solid lines show model fit, where each row represents a different model class, and data points show empirical data. Triangles data points are from highest intensity level and circles are from the lowest intensity level. Colours show proportions of category responses. For the audio-visual task, the predictions from best performing model in each class is shown. Errors bars show ± 1*SEM*.

#### Scaled Evidence Strength Models

According to the scaled evidence strength account, confidence reports depend on the distance between the observer’s internal representation of the stimulus and sensory uncertainty dependent choice and confidence criteria. We estimated values of *k* and *m* for each response criterion, which allowed the effect of sensory noise on the positioning of the criteria to differ across response options (k_r_ + m_r_*σ*^*z*^). This meant that the criteria associated with low confidence regions could expand under high sensory noise, while the corresponding high confidence regions could contract (and vice versa for low sensory noise). We also estimated an exponent on the measurement noise term, *z*, which was constant across criteria. The exponent allowed for the effect of *σ* on the choice and confidence criteria to change as a function of *σ*.

As shown in **Figure 7B**, the scaled evidence strength model was able to capture the empirical data well, where for the low intensity stimuli (those associated with higher sensory noise), confidence ratings remained uniformly low across stimulus values. For higher intensity stimuli (those associated with lower sensory noise), the criteria shifted so that higher confidence ratings were given for stimulus values associated with greater evidence strength (stimulus values with greater category diagnosticity) and lower confidence ratings were given for stimulus values associated with less evidence strength (stimulus values with lower category diagnosticity).

As shown in **Figure 8**, the scaled evidence strength models had the lowest summed *AIC* and *BIC* scores and they outperformed the alternative models for the majority of individual participants (see **Figure 9A**), especially for the unidimensional tasks. Specifically, the unscaled evidence strength models provided the best account for 8 out of the 10 participants in the auditory task and 9 out of 10 the participants in the visual task. We found that for the multidimensional task, the best performing model class was more heterogenous but that on average, the scaled evidence strength models still outperformed the other models, selected as the best performing class for 7 out of the 10 participants. The overall performance of the scaled evidence strength class suggests that observers consider sensory uncertainty when generating a confidence report, unlike the unscaled evidence strength models, but they do not do so in a way that is consistent with Bayesian inference (discussed in more detail below).

**Figure 8.**
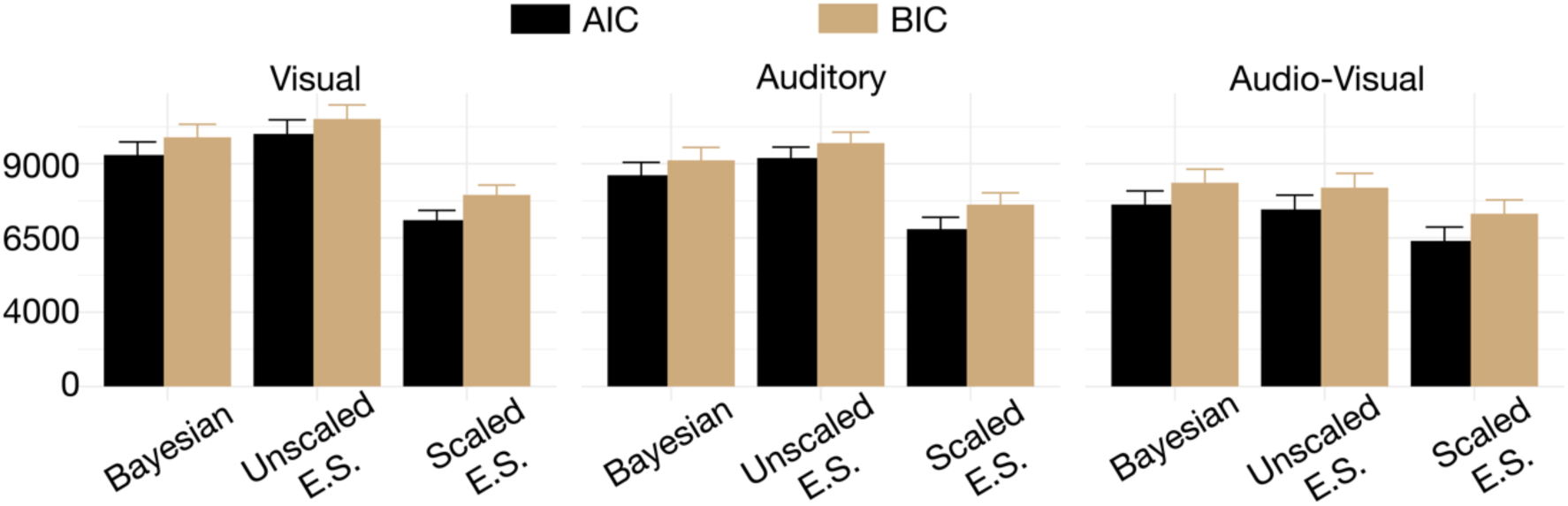
Summed AIC and BIC Across Participants For Each Model Class. *Note.* The scaled evidence strength class had the lowest summed scores across the visual (left panel), auditory (middle panel) and audio-visual (right panel) tasks.

#### Bayesian Models

According to Bayesian models of confidence, observers optimally combine sensory information with prior knowledge to form a posterior belief about the nature of the stimulus. In our task, we assumed that participants use the generating category distributions to calculate a log posterior ratio, 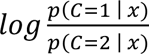 which they compare with choice-confidence criteria in the same measurement units. The Bayesian models have fewer parameters than the scaled evidence strength models because the choice and confidence criteria have the same positions in probability space but the estimated values of sensory noise at each intensity level allow these criteria to shift in perceptual space. That is, the amount of uncertainty associated with the sensory observation causes the associated response regions to expand or contract.

As shown in **Figure 7C**, the Bayesian models are not sufficiently flexible to capture the empirical confidence data. Specifically, while the model predictions are able to capture the empirical changes in responses as a function of stimulus values, where greater evidence strength is associated with greater confidence, the predictions show systematic deviations from the data across intensity levels. In particular, the Bayesian models consistently over-estimate confidence for stimuli of lower intensity and under-estimate confidence for stimuli of greater intensity. In terms of model selection, we found that the Bayesian models were selected by a single participant in the auditory and visual tasks and by 2 participants in the audio-visual task (see **Figure 9**). In terms of summed *AIC* and *BIC*, the Bayesian models were out-performed by the scaled evidence strength models (see **Figure 8**). The Bayesian models performed poorly even when we allowed for additional flexibility in the model by estimating participants’ prior knowledge of the generating category parameters, instantiating a subjective prior, rather than assuming they represent the true parameters veridically. Overall, these findings suggest that participants do not generate confidence reports in a way that is consistent with a Bayesian formalisation in any of our tasks.

**Figure 9.**
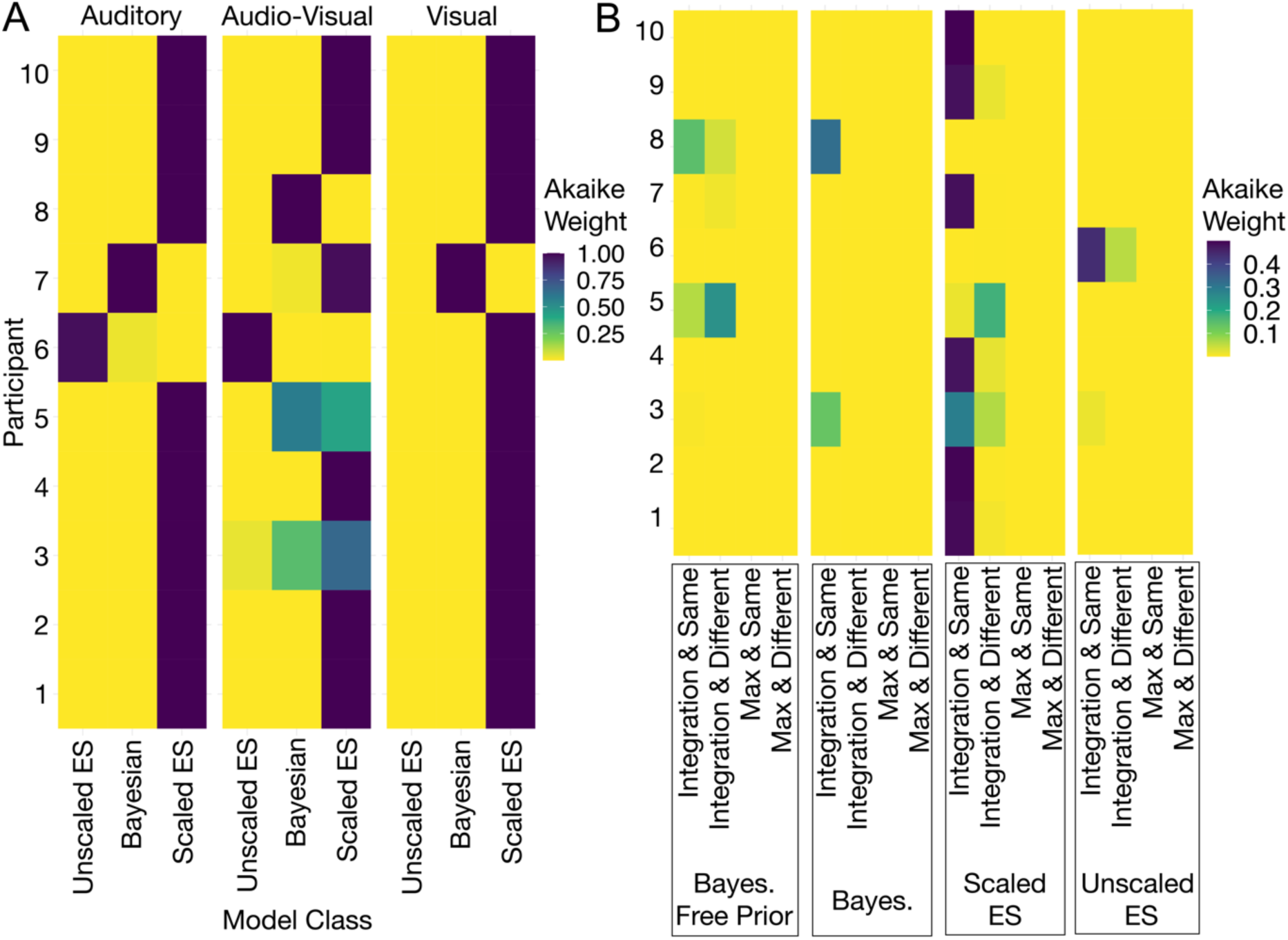
Akaike Weights Across Participants. *Note.* **(A)** Weights were calculated using the best-performing models in each class for each participant. Darker colours show larger weights, demonstrating the superior performance of the scaled evidence strength models across all tasks **(B)** Weights were calculated across all possible combinations of assumptions and model classes for the multidimensional task. Assumptions for model classes (Bayesian, scaled evidence strength and unscaled evidence strength), the combination of stimulus dimensions (integration or max) and sensory noise (same or different) are shown in different columns. Darker colours show larger weights, demonstrating the superior performance of the scaled evidence strength models with the integration and same noise assumption.

#### Predicting Multidimensional Behaviour from Unidimensional Behaviour

For the majority of cases, the scaled evidence strength models provided the best account of choice and confidence across modalities, consistent with the results from previous research using the same model classes (Adler & Ma, 2018; Denison et al., 2018; West et al., 2023). Given the similarities in the type of algorithm used across modalities, we wanted to investigate how well the confidence process generalised by investigating how well behaviour in the unidimensional tasks (visual and auditory) could predict behaviour in the multidimensional (audio-visual) task. To this end, we used the estimated parameters from each participant’s unidimensional model fits to predict their category and confidence responses to multidimensional stimuli. We did this using the model parameters from the scaled evidence strength model, because it provided the best fit across participants. We combined the parameters using the integration and different noise assumptions, as they consistently outperformed the max models and included the same noise as a special case.

Based on pilot data aimed at matching performance across tasks, we selected a more compressed range of intensity levels for the multidimensional task, where the second-highest intensity in the unidimensional tasks (level 3) corresponded to the highest intensity in the multidimensional task (level 4). To generate predictions for the multidimensional task, therefore, we estimated sensory noise parameters for the intermediate audio-visual intensity levels (intensity 2 and intensity 3), which were not tested in the unidimensional tasks. We fit an exponential function to the noise parameters from the four unidimensional intensity levels and used this function to predict noise values for the intermediate levels. The exponential function was chosen for convenience, as it enforced monotonicity and required estimating only a single parameter. We confirmed that it fit the data well (see **Figure S 2**). The tested (intensity 1 and 4) and interpolated (intensity 2 and 3) noise parameters provided noise estimates for each modality at the audio-visual intensity levels for each participant. These were then combined using Equation 12 to calculate *σ*_*AV*_.

We used the noise parameters to calculate choice and confidence criteria (*b*) in each modality, applying the estimated *k*, *m* and *z* parameters (see Equation 19) for each participant. These criteria were then transformed into a combined distance metric using the equation 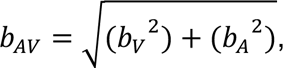 yielding predicted criteria, *b*_*AV*_, and noise, *σ*_*AV*_, parameters for each participant, directly estimated by the unidimensional model parameters.

In the final step, we estimated an attentional weight (*w*_*V*_) from the response and stimulus data in the multidimensional task. This weight was applied to the visual dimension, while the complementary weight (1 − *w*_*V*_) was applied to the auditory dimension. By applying this weighting, we captured systematic biases in participants’ reliance on one stimulus dimension over the other. We then calculated the likelihood of the dataset given the set of model parameters (see Equation 23). This model, which generalises parameters from the unidimensional tasks to the multidimensional task, is referred to as the *dimensionality-independent* model.

We found that with a single free parameter (the attentional weight), the dimensionality-independent model predictions captured the empirical data from the multidimensional task well (see **Figure 10**). The model did, however, underestimate confidence for stimulus values associated with greater category diagnosticity (values from the middle of the category 1 distribution and from the tails of the category 2 distribution), highlighting areas where confidence judgements may require further refinement. Overall, however, the dimensionality-independent model demonstrated strong predictive power across tasks, providing further evidence for the generalisability of a single confidence algorithm across different types of perceptual decisions.

**Figure 10.**
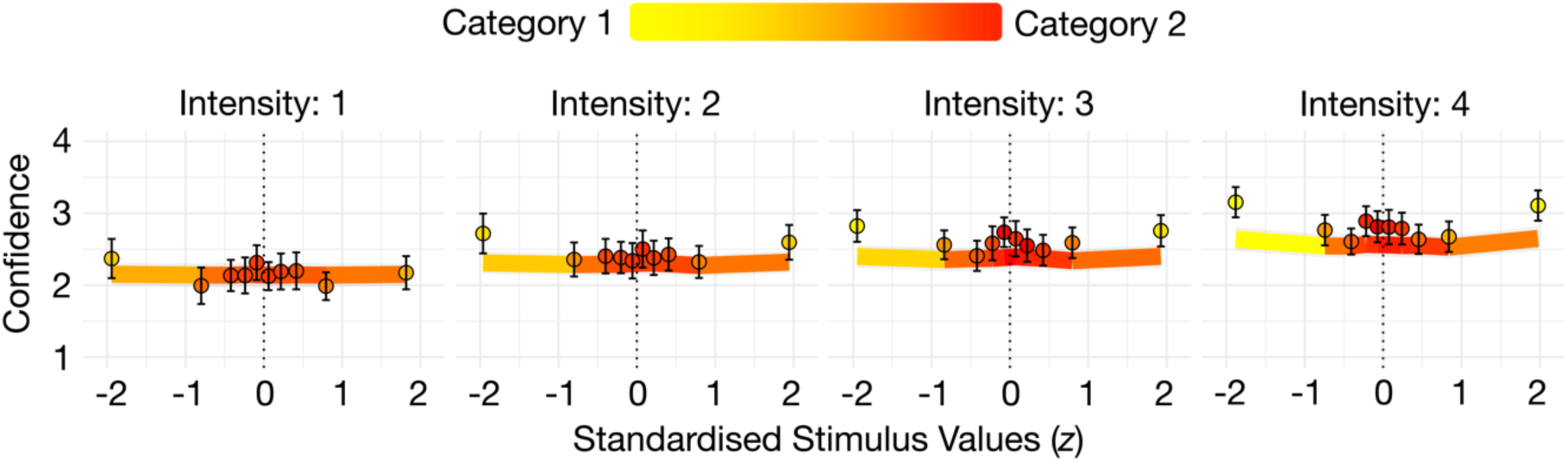
Dimensionality-Independent Model Predictions. *Note.* Model predictions from the dimensionality-independent model in which we used the estimated parameters from the unidimensional (visual and auditory) model fits to predict responses on the multidimensional (audio-visual) task. Solid lines show model predictions and circular plotting symbols show empirical data from the audio-visual task. Colours show proportion of category responses. Error bars show ± 1 *SEM*.

#### How is Sensory Information Combined Across Modalities?

Relatively little research has explored how confidence models perform in tasks requiring the integration of information across sensory modalities. In this study, therefore, we wanted to use our modelling approach to more comprehensively characterise choice and confidence judgements in the audio-visual task. To do so, we tested two key assumptions about the combination of sensory information across modalities for each of the model classes, resulting in 4 variants for each core model (i.e., a total of 16 models were fit to the audio-visual data across the 3 model classes).

The first set of assumptions allowed us to investigate whether participants attended to both dimensions of the multidimensional stimulus. In the ‘integration’ models, we assumed that participants attended to both dimensions, while in the ‘max’ models, we assumed that they attended only to the most informative dimension. We found that the integration models outperformed the max models in all classes for every participant, confirming that confidence judgements relied on combining auditory and visual input. However, participants did not appear to integrate stimulus values across dimensions in an unbiased way. Parameter estimates showed that all participants were biased towards the visual modality, weighting the visual dimension of the stimulus more heavily in their confidence judgement.

The second set of assumptions explored how participants represented sensory noise. We implemented two model instantiations, one in which we estimated noise values separately for each stimulus dimension (these estimates were later combined across the two modalities) and the other in which we estimated a single ‘modality-independent’ sensory noise value. Models with a modality-independent noise parameter performed better, suggesting participants rely on a generalised noise estimate rather than combining noise from individual dimensions.

Together, these findings help elucidate how participants generate confidence reports when integrating sensory information. We showed that confidence judgements were based on both stimulus dimensions, instantiated in the model as a weighted ‘distance’ measure. This combined measure, which provided a proxy for evidence strength, was compared against choice and confidence criteria that were adjusted based on a modality-independent noise estimate which varied with stimulus intensity. As shown in **Figure 7B**, this model closely aligned with empirical data, capturing response variation driven by stimulus values (evidence strength) and intensity (measurement noise).

## Discussion

Most previous studies of the computational basis of decision confidence have relied on simple visual perceptual tasks, leaving unanswered how confidence is formed in decisions involving other sensory modalities or those that require integrating information across modalities. To address these questions, we used computational modelling to analyse confidence judgements in perceptual decision-making tasks involving visual, auditory and audio-visual stimuli. Our first set of analyses investigated whether confidence judgements for such decisions are based on the same underlying computational algorithms. Specifically, we compared the performance of different classes of confidence models and evaluated whether the same model class consistently fit data from all three tasks. We further tested the generalisability of a single class of algorithms by examining how well predictions generated from the unidimensional tasks (visual and auditory) aligned with empirical data from the multidimensional (audio-visual) task. The second set of analyses focused specifically on the multidimensional task, examining how participants integrated sensory inputs from multiple modalities when making confidence judgements. Within each model class, we tested different assumptions about the integration process. In the sections that follow, we first discuss the model classes and their performance across the different tasks, and then provide a more detailed analysis of the modelling results from the multidimensional task.

### A Theoretical Framework for Understanding Confidence Judgements: Comparing Model Classes

Building on an extensive body of research on the computational basis of confidence in visual perceptual tasks (Adler & Ma, 2018; Aitchison et al., 2015; Faivre et al., 2018; Locke et al., 2022; Navajas et al., 2017; Shekhar & Rahnev, 2024; West et al., 2023), we adapted models to evaluate how well they fit the data from our visual, auditory and novel audio-visual categorisation tasks. Our model classes instantiated different assumptions about how observers transform sensory signals, whether unidimensional or multidimensional, into a choice and corresponding confidence report. Within our general modelling framework, we assumed that sensory input from the external world generates an internal response, represented as a latent decision variable. The latent variable was modelled as a normal distribution, to capture the noise inherent in the sensory system itself. A key aspect of all our models, therefore, was the concept of *sensory uncertainty*, which refers to the variability (or noise) in the observer’s internal representation of the stimulus. We assumed that this sensory uncertainty (characterised by the *σ* parameters) varied with stimulus intensity, with lower intensities leading to greater uncertainty. Sensory uncertainty also influenced the choice and confidence criteria, and different model classes handled this relationship in distinct ways.

For the non-Bayesian models, the decision variable was expressed in psychological units which were tied directly to the stimulus (e.g., a grating’s orientation). The value of the stimulus itself, therefore, guided both category choice and confidence, with the choice-confidence criteria segmenting the possible values into discrete category and confidence response regions. We tested two types of non-Bayesian models: one which assumed fixed choice-confidence-criteria regardless of sensory uncertainty (unscaled evidence strength; Kepecs et al., 2008; Komura et al., 2013; Rahnev et al., 2011), and the other which assumed that the choice-confidence-criteria were adjusted based on the amount of sensory uncertainty in the decision variable (scaled evidence strength; Adler & Ma, 2018; Denison et al., 2018; Locke et al., 2022; West et al., 2023). In contrast, the Bayesian models approached sensory uncertainty by incorporating it into a probabilistic framework. These models used prior knowledge about the generating category distributions to compute a log posterior probability ratio. The magnitude of the log posterior probability ratio served as a choice and confidence metric, directly reflecting how sensory uncertainty impacts the perceived likelihood of a decision outcome.

#### Bayesian Class

We found that that Bayesian model class was outperformed across all tasks, both at the group level and for the majority of individual participants. Even after increasing their flexibility, by estimating participants’ subjective prior knowledge rather than assuming they knew the true generating category parameters, the performance of the Bayesian models remained relatively poor. This consistently poor performance suggests that participants’ confidence judgements in all our tasks are unlikely to rely on posterior probability computations, at least as represented by the models we tested. These findings align with previous research demonstrating the limited applicability of Bayesian models to metacognitive decisions (Adler & Ma, 2018; Baranski & Petrusic, 1994; Locke et al., 2022; West et al., 2023). Notably, our results extend previous research by showing that confidence does not follow a strict Bayesian framework, even in multidimensional contexts where a modality-independent measurement system could theoretically resolve differences in how information from distinct sensory modalities is represented and integrated, resolving any ‘translation issues’. The underperformance of the Bayesian models, even in tasks where integration may benefit from a unified measurement system, highlights the broader limitations of these models. Specifically, the rigid assumptions inherent to Bayesian inference may fail to account for the context-dependent nature of human confidence judgements.

Additionally, these findings suggest that the use of prior information in confidence judgements may be less straightforward than previously assumed. Rather than incorporating priors in a statistically optimal manner, individuals may employ them more heuristically, in ways that do not adhere to the strict probabilistic structure imposed by Bayesian models. This highlights the importance of future research exploring alternative modelling frameworks that incorporate prior knowledge in a more flexible and computationally efficient manner.

#### Unscaled Evidence Strength Class

In contrast to the Bayesian models, the unscaled evidence strength models instantiated the simplest view of metacognitive computations. In the unscaled evidence strength models, the decision variable, which was directly related to the stimulus itself, was compared to choice and confidence criteria that did not change as a function of sensory uncertainty. The unscaled evidence strength model performed worst on all tasks, except in a small number of individual cases (one participant in the auditory task and one participant in the audio-visual task). This finding aligns with previous studies that have documented poor performance of unscaled evidence strength models across unidimensional tasks performed in vision (Adler & Ma, 2018) and audition (West et al., 2023). The results from our study extend this understanding to multidimensional decisions, suggesting that even when observers must integrate stimulus information from distinct modalities, they rely on more sophisticated computations that factor in sensory uncertainty.

#### Scaled Evidence Strength Class

In the scaled evidence strength models, we assumed that the decision variable was directly related to the value of the stimulus but that the choice and confidence criteria were dynamically adjusted in response to sensory uncertainty. This approach incorporated a scaling factor (*mσ*^*z*^) that adjusted the criteria, accommodating linear, quadratic and faster than quadratic relationships between sensory uncertainty and criterion positioning. This flexibility allowed the model to capture individual differences in how participants adapted to varying levels of noise. The scaled evidence strength models outperformed all alternative models across tasks, both for the majority of individual participants and for the group.

The scaled evidence strength framework allowed observers to integrate sensory uncertainty into their confidence judgements, much like an ‘optimal observer,’ but without the computational burden of posterior probability computations. The success of the scaled evidence strength class aligns with emerging evidence that confidence judgements rely on heuristic-like strategies that approximate optimal behaviour (Adler & Ma, 2018; Aitchison et al., 2015; Denison et al., 2018; West et al., 2023), extending these findings to decisions involving the integration of multiple sources of information. The superior performance of this class of models suggests that future studies may benefit from including models that incorporate scaling mechanisms which allow confidence to dynamically adapt to uncertainty, without relying on posterior probabilities.

#### A Shared Computational Basis for Confidence

The finding that the scaled evidence strength models outperformed the other models across all tasks supports the idea that the same core algorithm governs confidence judgements across modalities. We further investigated the generalisability of the confidence process by testing whether behaviour in the unidimensional tasks (visual and auditory) could predict behaviour in the multidimensional (audio-visual) task using the scaled evidence strength algorithm. Specifically, we used the parameters estimated from the unidimensional model fits for each participant and estimated a single additional parameter to predict their category and confidence responses to multidimensional stimuli. This approach, referred to as the dimensionality-independent model, revealed that predictions generated from the unidimensional tasks closely aligned with the observed data from the audio-visual task. This finding reinforces the notion of a shared algorithmic basis for confidence judgements across different types of perceptual decisions and aligns with both our previous computational work (West et al., 2023) and other related studies (Ais et al., 2016; Faivre et al., 2018; Fleck et al., 2006; Heereman et al., 2015; Mazancieux et al., 2020; Morales et al., 2018; Pleskac & Busemeyer, 2010; Rouault et al., 2018; Song et al., 2011) showing common mechanisms for confidence judgements across modalities.

It is important to acknowledge, however, that our conclusions about a shared algorithmic basis are limited by the scope of the models we tested. While we included a range of models grounded in existing theoretical frameworks, it remains possible that alternative models, either Bayesian or non-Bayesian, might account for the data equally well or better. Nevertheless, the fact that the scaled evidence strength algorithm outperformed other alternatives across all tasks, along with its close fit to the data in each case, suggests that it captures a core feature of how humans generate confidence judgements.

Looking ahead, future studies could explore the use of latent factor models to gain deeper insights into the underlying shared computational mechanisms (Schmiedek et al., 2007; Stevenson et al., 2024). In the context of our study, a latent factor model could capture underlying cognitive constructs – such as modality-independent confidence mechanisms – that operate across different sensory tasks. Specifically, instead of modelling each task separately, a latent factor model could account for both task-specific parameters and shared latent constructs that generalise across tasks. By modelling both the latent cognitive constructs and task parameters simultaneously, such models could provide insights into whether the underlying confidence algorithm truly operates independently of the sensory modality or if some modality-specific factors still play a role.

### Metacognition Across Stimulus Dimensions: Comparing Assumptions for Integrating Information Across Modalities

To the best of our knowledge, few studies have used computational modelling to characterise the computations involved in metacognitive judgements when integrating stimulus information across sensory modalities. Addressing this gap, we developed a set of models to explore how people form confidence judgements when integrating information from visual and auditory sources. In doing so, we examined two assumptions about the confidence generation process.

The first set of assumptions quantified whether participants integrated both stimulus dimensions or whether they relied on a single dimension only. In the *integration* assumption, the decision variable was derived from a weighted integration of both dimensions of the stimulus, whereas in the *max* assumption, it relied solely on the dimension providing the strongest input. Our results showed consistent evidence in favour of the integration assumption, with integration models outperforming the max models across all classes and participants. This finding aligns with prior research on perceptual decisions, which showed that people are able to integrate information from different sensory modalities (Alais & Burr, 2004; Calvert et al., 1998; Driver & Spence, 1998; Ernst & Banks, 2002; McGurk & MacDonald, 1967; Shams et al., 2000; Stein & Meredith, 1993; Van Wassenhove et al., 2005).

This result also aligns with preliminary work that has investigated metacognition for perceptual decisions involving multiple modalities (Faivre et al., 2018; Gao et al., 2023). For example, Gao and colleagues (2023) demonstrated that sensory cues are automatically integrated based on their reliability, even when participants were instructed to disregard one of the modalities because it was task-irrelevant. Our results build on those from Gao and colleagues (2023) by showing that a reliability-weighted integration process also operates when both modalities are explicitly task-relevant. This finding supports the idea that the integration process reflects a broader, more fundamental principle of information processing. Importantly, it shows that integration is not merely an artefact of a failure in selective attention; rather, the brain appears to automatically combine information from all available modalities, even if that information is not directly task-relevant.

Faivre and colleagues (2018) also explored confidence in audio-visual decisions, demonstrating through computational modelling that confidence could be predicted either from a bidimensional representation of an audio-visual signal or by comparing unidimensional visual and auditory representations. Our approach extends the work of Faivre and colleagues (2018) by considering a broader range of theoretical model classes beyond modelling confidence as the probability of being correct, an assumption that aligns most closely with our Bayesian models. This broader modelling framework allowed us to investigate multiple possible algorithms underlying confidence generation, providing deeper insight into how sensory information is integrated at the metacognitive level.

Specifically, the weight parameter in our integration models also allowed us to directly investigate whether participants systematically over-weighted one stimulus dimension in their confidence judgement. All participants exhibited a bias toward the visual modality. This over-reliance on visual information aligns with previous studies suggesting that visual input tends to dominate multisensory integration (Alais & Burr, 2004; Colavita, 1974; Ernst & Banks, 2002; McGurk & MacDonald, 1967; Posner et al., 1976; Rock & Victor, 1964; Welch & Warren, 1980). Importantly, our results demonstrate that this visual dominance extends beyond perceptual judgements to influence confidence judgements as well, reinforcing the idea that confidence reflects the same biases present in perception. Further research could examine the relative weighting of visual and auditory inputs more closely. One possible avenue would be to implement dynamic models that update weighting on a trial-by-trial basis, depending on stimulus characteristics. For example, the model could reflect a baseline bias toward the visual modality that becomes less dominant when auditory cues provide highly informative information. Another potential direction would involve trying to experimentally manipulate the integration weights by instructing participants to attend selectively to specific stimulus dimensions. This would allow future studies to also explore whether these biases are modifiable through top-down attentional control.

The second set of assumptions that we examined in our multidimensional models was how participants estimated sensory uncertainty across modalities. We tested two model variants to address this question: one in which sensory noise values were estimated separately for each stimulus dimension and then combined across modalities, and another in which a single modality-independent sensory noise value was used across both dimensions. Our results indicated that participants adjusted their choice and confidence criteria based on a single estimate of sensory uncertainty, which varied systematically with the experimentally manipulated intensity. However, this result should be interpreted cautiously, as our model recovery analysis indicated a slight over-selection of models with the same noise parameters across modalities, potentially biasing this conclusion (see **Figure 5**).

Despite this, we propose several possible interpretations to guide future research. One is that the noise estimate in the multidimensional models may serve as a general measure of uncertainty, rather than capturing low-level sensory noise specific to each modality (e.g., noise which could be represented as variance in neuron firing rates). From this perspective, participants may not require a fine-grained noise estimate for each sensory input because a single general uncertainty measure suffices to regulate confidence across modalities. This interpretation aligns with other computational models of confidence including the weighted evidence and visibility (WEV) model (Rausch et al., 2018, 2021). The WEV model posits that individuals use visibility as a global signal shaping confidence judgements, regardless of the specific sensory modality contributing to the evidence. This visibility signal functions similarly to a global noise estimate, modulating the level of confidence placed in the accumulated evidence.

Alternatively, it is possible that our intensity manipulation was effective in matching the amount of sensory uncertainty across modalities, making a single noise estimate sufficient to capture the amount of low-level sensory noise in each modality. However, this explanation seems unlikely given the observed differences in categorisation performance at different intensity levels in the unidimensional tasks (see **Figure S 1**). An interesting direction for future research would be to vary intensity independently within each stimulus dimension, e.g., using high-intensity visual input combined with low-intensity auditory input. This approach would allow for a more nuanced investigation of whether participants can dynamically adjust their confidence judgements based on independent sensory noise levels across modalities.

### Conclusions

In summary, our findings demonstrate that the scaled evidence strength models consistently outperformed the other model classes across all tasks and could be used to predict behaviour in the multidimensional task based on unidimensional data. These findings suggest that confidence judgements are governed by a shared algorithm that dynamically accounts for both sensory uncertainty and evidence strength, without relying on posterior probability computations. Additionally, we demonstrated that when making multidimensional confidence judgements, participants integrated visual and auditory information with a systematic bias toward the visual modality, and used a modality-independent measure of sensory uncertainty to modulate their confidence. Our results highlight the generalisability of the scaled evidence strength algorithm, even for tasks requiring integration across sensory modalities, and emphasise the limitations of Bayesian models in capturing the nuanced, context-dependent nature of human confidence judgements. These results open important avenues for future research, including the use of latent factor models to capture both shared and task-specific metacognitive constructs, exploring how sensory inputs are dynamically weighted for confidence, and investigating whether integration biases in confidence can be modulated through top-down attentional control.

## Supporting information

All Supplemental

