## Supplementary material for "A common algorithm for confidence judgements across visual, auditory and audio-visual decisions": All Supplemental

### Supplementary Materials

**Figure S 1**

*Categorisation Accuracy Across Stimulus Intensities*

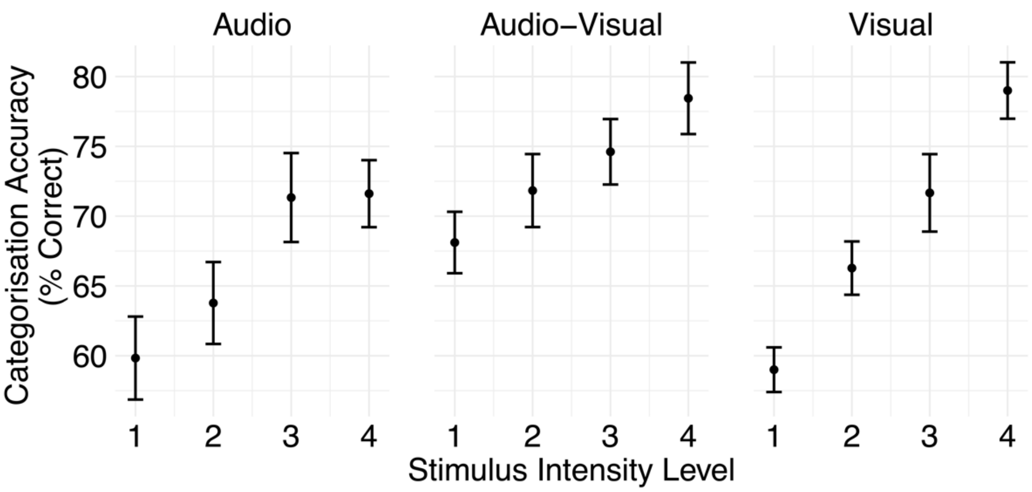


*Note.* Error bars show $\pm 1\, SEM$.

**Table S 1**

*Categorisation Accuracy for Highest Intensity Stimuli*

| Participant | Visual | Auditory | Audio-Visual |
|---|---|---|---|
| 1 | 87.78 | 78.89 | 86.67 |
| 2 | 82.22 | 70.00 | 77.78 |
| 3 | 75.56 | 81.11 | 84.44 |
| 4 | 72.78 | 77.78 | 81.11 |
| 5 | 82.22 | 81.11 | 86.11 |
| 6 | 67.78 | 62.78 | 67.22 |
| 7 | 76.11 | 66.11 | 70.56 |
| 8 | 75.56 | 61.11 | 78.33 |
| 9 | 85.56 | 66.67 | 65.56 |
| 10 | 84.44 | 70.56 | 86.67 |

*Note*. To ensure that all participants had a clear understanding of the task and underlying category distributions, we assessed categorisation accuracy for each participant using a binomial significance test. This test compared each participant's accuracy in each task to chance performance, using data from testing trials in the highest intensity condition only. Categorisation accuracy was calculated based on the most probable generating category rather than the true generating category.

#### Figure S 2

*Noise Parameters Across Stimulus Intensities*

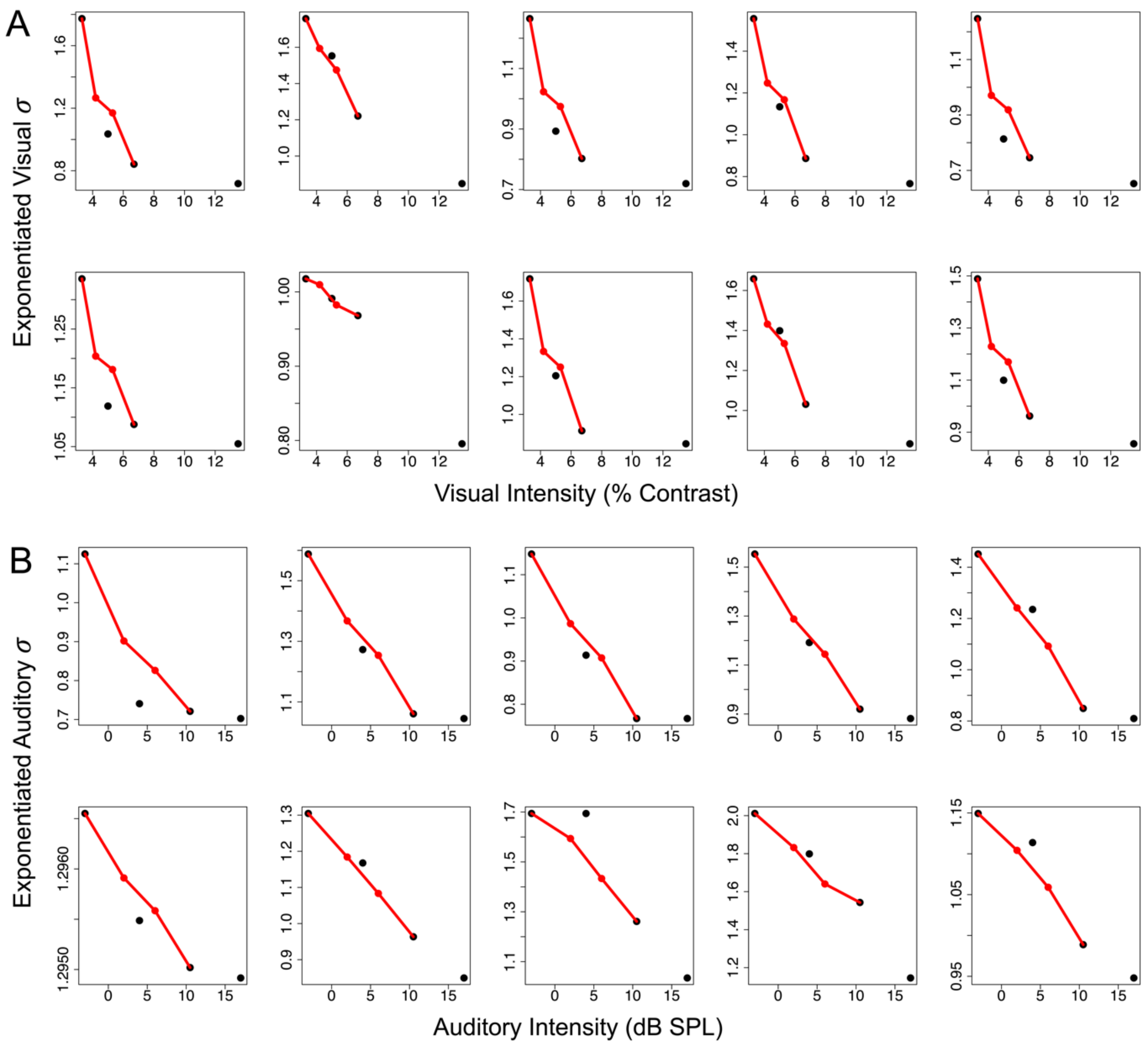


*Note.* Black points show estimated noise parameters at the tested intensity levels in the unidimensional tasks for **(A)** visual and **(B)** auditory modality. An exponential function was fit to the estimated parameters from the unidimensional tasks. Red points show predicted noise parameters at intermittent bidimensional intensity levels.

**Table S 2**

*Model-Free GLMMs for Category and Confidence Responses: Follow Ups*

| | | Tasks | | |
|---|---|---|---|---|
| | | Visual | Auditory | Audio-Visual |
| | | Category | | |
| Intensity 1 | OR | 1.22 | 1.25 | 2.53 |
| | $CI_{lower}$ | 1.03 | 0.87 | 2.00 |
| | $CI_{upper}$ | 1.45 | 1.80 | 3.19 |
| | *p* | 0.019 | 0.219 | < 0.001 |
| Intensity 2 | OR | 1.97 | 1.91 | 2.93 |
| | $CI_{lower}$ | 1.61 | 1.26 | 2.21 |
| | $CI_{upper}$ | 2.42 | 2.92 | 3.90 |
| | *p* | < 0.001 | 0.003 | < 0.001 |
| Intensity 3 | OR | 3.12 | 3.27 | 3.73 |
| | $CI_{lower}$ | 2.19 | 2.04 | 2.82 |
| | $CI_{upper}$ | 4.46 | 5.23 | 4.92 |
| | *p* | < 0.001 | < 0.001 | < 0.001 |
| Intensity 4 | OR | 5.88 | 3.62 | 4.66 |
| | $CI_{lower}$ | 4.41 | 2.50 | 3.19 |
| | $CI_{upper}$ | 7.83 | 5.23 | 6.82 |
| | *p* | < 0.001 | < 0.001 | < 0.001 |
| | | Confidence | | |
| Intensity 1 | $\beta$ | 0.02 | 0.01 | 0.06 |
| | $CI_{lower}$ | -0.02 | -0.03 | 0.01 |
| | $CI_{upper}$ | 0.06 | 0.05 | 0.10 |
| | *p* | 0.244 | 0.643 | 0.015 |
| Intensity 2 | $\beta$ | 0.04 | 0.12 | 0.05 |
| | $CI_{lower}$ | -0.01 | 0.02 | -0.02 |
| | $CI_{upper}$ | 0.08 | 0.21 | 0.12 |
| | *p* | 0.097 | 0.016 | 0.152 |
| Intensity 3 | $\beta$ | 0.10 | 0.12 | 0.09 |
| | $CI_{lower}$ | 0.04 | -0.03 | 0.02 |
| | $CI_{upper}$ | 0.16 | 0.26 | 0.16 |
| | *p* | 0.001 | 0.108 | 0.012 |
| Intensity 4 | $\beta$ | 0.18 | 0.11 | 0.13 |
| | $CI_{lower}$ | 0.10 | -0.03 | 0.04 |
| | $CI_{upper}$ | 0.25 | 0.24 | 0.22 |
| | *p* | < 0.001 | 0.118 | 0.004 |

*Note.* CI terms refer to 95% confidence intervals calculated using the profile likelihood method. Significance values are obtained using the Satterthwaite approximation to calculate the degrees of freedom for the t-distribution based on the estimated variance-covariance matrix of the model parameters (Lüdecke, 2022). For category GLMMs (top section), we report the odds ratio (exponentiated coefficient estimate) for ease of interpretation. The odds ratio represents the change in odds of the outcome for a one-unit change in the predictor variable. For confidence GLMMs (bottom section), we report standardised regression coefficients.

**Table S 3**

*Parameter Recovery for Unscaled Evidence Strength Model Variants*

| | Max – Same Noise | Integrated – Same Noise | Max – Different Noise | Integrated – Different Noise |
|---|---|---|---|---|
| $b_1$ | 0.97 *** | 1.00 *** | 1.00 *** | 1.00 *** |
| $b_2$ | 0.98 *** | 0.99 *** | 0.99 *** | 1.00 *** |
| $b_3$ | 0.99 *** | 0.98 *** | 0.98 *** | 1.00 *** |
| $b_4$ | 1.00 *** | 0.98 *** | 0.99 *** | 1.00 *** |
| $b_5$ | 0.99 *** | 0.97 *** | 0.95 *** | 1.00 *** |
| $b_6$ | 0.97 *** | 0.99 *** | 0.91 *** | 0.99 *** |
| $b_7$ | 0.98 *** | 0.97 *** | 0.93 *** | 0.99 *** |
| $\sigma_{V[1]}{}^{\circ}$ | 0.92 *** | 0.78 ** | 0.90 *** | 0.09 |
| $\sigma_{V[2]}{}^{\circ}$ | 0.96 *** | 0.82 ** | 0.95 *** | 0.34 |
| $\sigma_{V[3]}{}^{\circ}$ | 0.99 *** | 0.94 *** | 0.99 *** | 0.71 * |
| $\sigma_{V[4]}{}^{\circ}$ | 0.99 *** | 0.99 *** | 0.97 *** | 0.29 |
| $w$ | | 0.95 *** | | 0.88 *** |
| $\sigma_{A[1]}$ | | | 0.91 *** | 0.45 |
| $\sigma_{A[2]}$ | | | 0.96 *** | 0.46 |
| $\sigma_{A[3]}$ | | | 0.97 *** | 0.32 |
| $\sigma_{A[4]}$ | | | 0.72 * | 0.69 * |

*Note*: *$p$ < 0.05, **$p$ < 0.01, ***$p$ < .001. °For models with the same noise assumption, $\sigma$ parameters are modality independent but are shown in rows labelled $\sigma_V$.

**Table S 4**

*Parameter Recovery for Scaled Evidence Strength Model Variants*

| | Max – Same Noise | Integrated – Same Noise | Max – Different Noise | Integrated – Different Noise |
|---|---|---|---|---|
| exponent | 0.79 ** | 0.95 *** | 0.96 ** | 0.86 ** |
| $k_1$ | 0.99 *** | 1.00 *** | 0.99 *** | 0.96 *** |
| $k_2$ | 0.98 *** | 0.98 *** | 0.99 *** | 0.99 *** |
| $k_3$ | 0.96 *** | 0.98 *** | 0.99 *** | 0.98 *** |
| $k_4$ | 0.95 *** | 0.84 ** | 0.98 *** | 0.98 *** |
| $k_5$ | 0.88 *** | 0.92 *** | 0.96 *** | 0.96 *** |
| $k_6$ | 0.70 * | 0.94 *** | 0.92 *** | 0.82 ** |
| $k_7$ | 0.65 * | 0.80 ** | 0.73 * | 0.95 *** |
| $m_1$ | 0.96 *** | 0.99 *** | 0.99 *** | 0.98 *** |
| $m_2$ | 0.94 *** | 0.98 *** | 1.00 *** | 0.98 *** |
| $m_3$ | 0.79 ** | 0.94 *** | 0.99 *** | 0.98 *** |
| $m_4$ | 0.99 *** | 0.94 *** | 0.96 *** | 0.98 *** |
| $m_5$ | 0.53 | 0.80 ** | 0.92 *** | 0.97 *** |
| $m_6$ | 0.47 | 0.96 *** | 0.92 *** | 0.93 *** |
| $m_7$ | 0.64 * | 0.81 ** | 0.76 ** | 0.89 ** |
| $\sigma_{V[1]}$° | 0.90 *** | 0.99 *** | 0.99 *** | 0.91 *** |
| $\sigma_{V[2]}$° | 0.96 *** | 0.98 *** | 0.96 *** | 0.92 *** |
| $\sigma_{V[3]}$° | 0.97 *** | 0.90 *** | 0.98 *** | 0.97 *** |
| $\sigma_{V[4]}$° | 0.96 *** | 0.91 *** | 0.99 *** | 0.95 *** |
| $w$ | | 0.77 ** | | 0.91 *** |
| $\sigma_{A[1]}$ | | | 0.83 *** | 0.95 *** |
| $\sigma_{A[2]}$ | | | 0.94 *** | 0.92 *** |
| $\sigma_{A[3]}$ | | | 0.97 *** | 0.85 ** |
| $\sigma_{A[4]}$ | | | 0.97 *** | 0.74 * |

*Note*: **p* < 0.05, ***p* < 0.01, ****p* < .001. °For models with the same noise assumption, $\sigma$ parameters are modality independent but are shown in rows labelled $\sigma_V$.

**Table S 5**

*Parameter Recovery for Bayesian Model Variants*

| | Max – Same Noise | Integrated – Same Noise | Max – Different Noise | Integrated – Different Noise |
|---|---|---|---|---|
| $b_1$ | 0.99 *** | 0.98 *** | 1.00 *** | 0.96 *** |
| $b_2$ | 0.96 *** | 0.99 *** | 0.98 *** | 0.97 *** |
| $b_3$ | 0.98 *** | 0.97 *** | 0.97 *** | 0.99 *** |
| $b_4$ | 0.97 *** | 0.97 *** | 0.97 *** | 0.97 *** |
| $b_5$ | 0.93 *** | 0.99 *** | 0.99 *** | 0.95 *** |
| $b_6$ | 0.97 *** | 0.98 *** | 0.65 * | 0.84 ** |
| $b_7$ | 0.88 ** | 0.92 *** | 0.97 *** | 0.98 *** |
| $\sigma_{V[1]}{}^{\circ}$ | 0.77 ** | 0.80 ** | 0.88 ** | 0.91 *** |
| $\sigma_{V[2]}{}^{\circ}$ | 0.97 *** | 0.81 ** | 0.85 ** | 0.83 ** |
| $\sigma_{V[3]}{}^{\circ}$ | 0.98 *** | 0.72 * | 0.91 *** | 0.73 * |
| $\sigma_{V[4]}{}^{\circ}$ | 0.99 *** | 0.97 *** | 0.91 *** | 0.79 ** |
| $w$ | | 0.95 *** | | 0.98 *** |
| $\sigma_{A[1]}$ | | | 0.88 ** | 0.35 |
| $\sigma_{A[2]}$ | | | 0.83 ** | -0.06 |
| $\sigma_{A[3]}$ | | | 0.93 *** | 0.29 |
| $\sigma_{A[4]}$ | | | 0.97 *** | 0.58 |

*Note*: *$p < 0.05$, **$p < 0.01$, ***$p < .001$. °For models with the same noise assumption, $\sigma$ parameters are modality independent but are shown in rows labelled $\sigma_V$.

**Figure S 3**

*Parameter Recovery for Unscaled Evidence Strength Models*

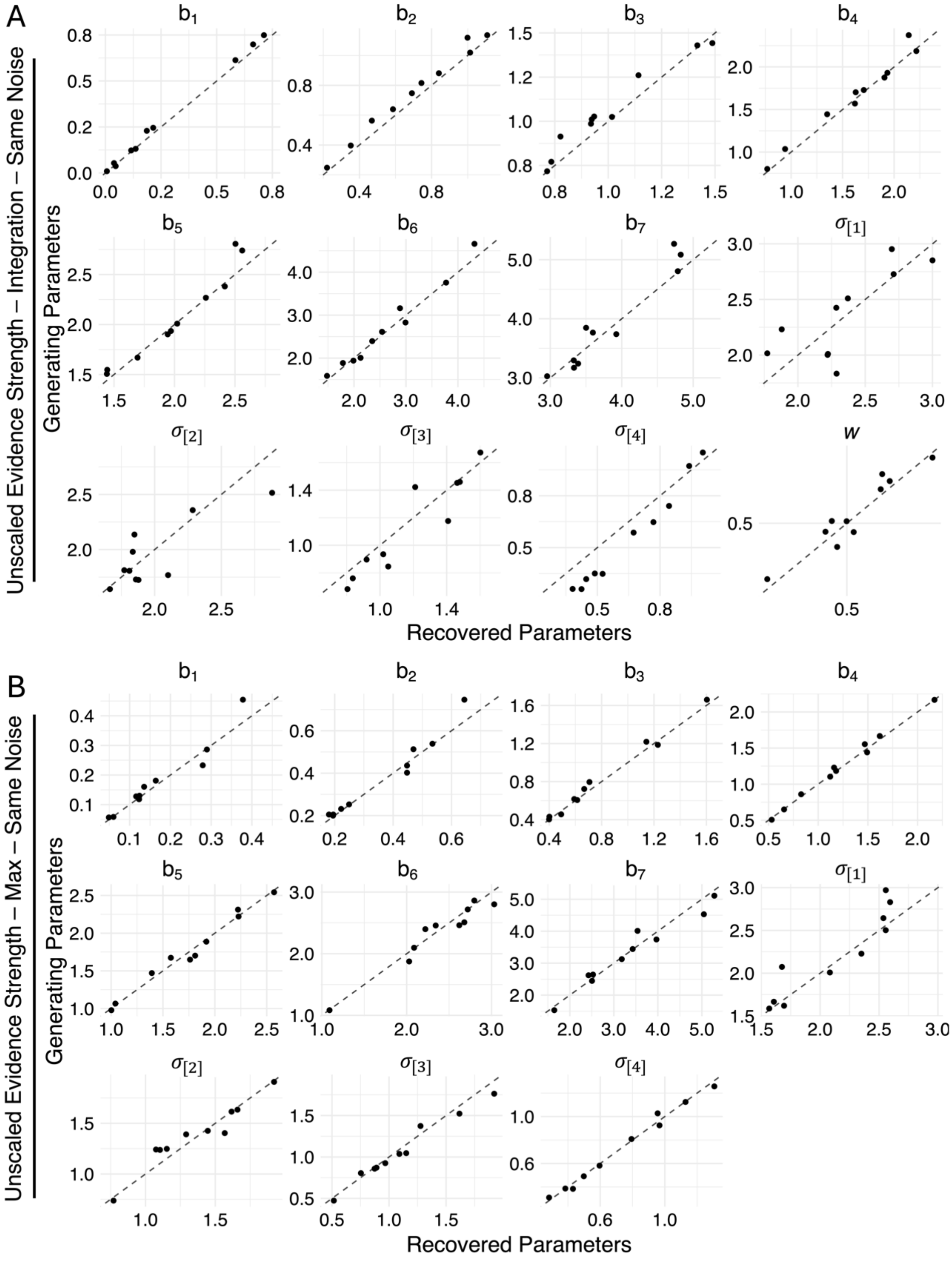


Recovery results for **(A)** integration same noise variant, **(B)** max same noise variant, (C) integration different noise variant, and (D) max different noise variant**.** $b_{1-7}$ refer to the category/confidence boundaries and $\sigma_{1-4}$ refer to the estimated noise parameters at each intensity level (the perceived level of sensory uncertainty in the observer's perception of the stimulus). *w* refers to the weight applied to the visual dimension during integration, where the auditory weight is equal to 1 – *w*. Dotted diagonal lines indicate perfect recovery of the data-generating parameter values.

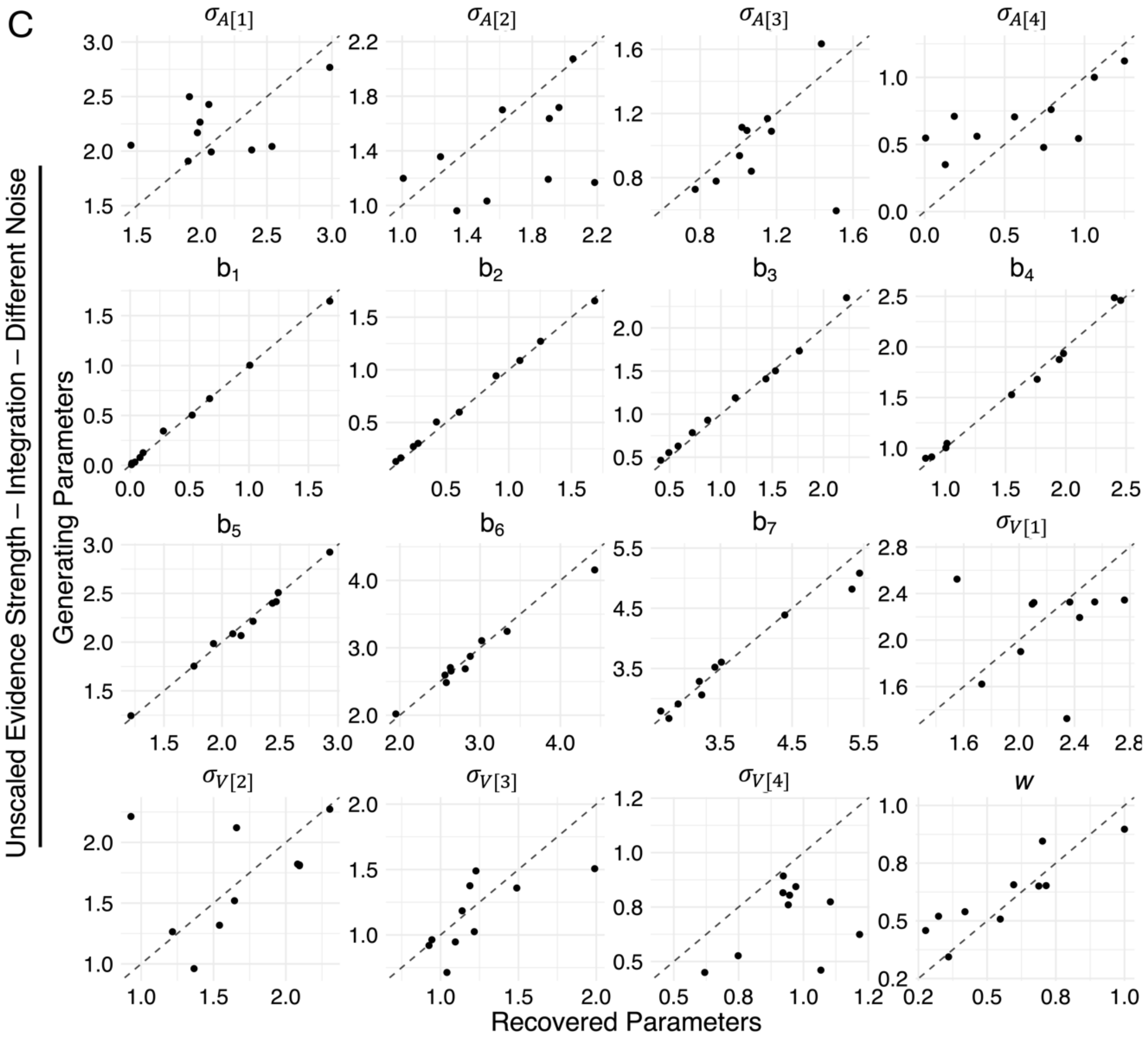


*Note.* Recovery results for (A) integration same noise variant, (B) max same noise variant, **(C)** integration different noise variant, and (D) max different noise variant**.** $b_{1-7}$ refer to the category/confidence boundaries and $\sigma_{1-4}$ refer to the estimated noise parameters at each intensity level (the perceived level of sensory uncertainty in the observer's perception of the stimulus). *w* refers to the weight applied to the visual dimension during integration, where the auditory weight is equal to 1 – *w*. Dotted diagonal lines indicate perfect recovery of the data-generating parameter values.

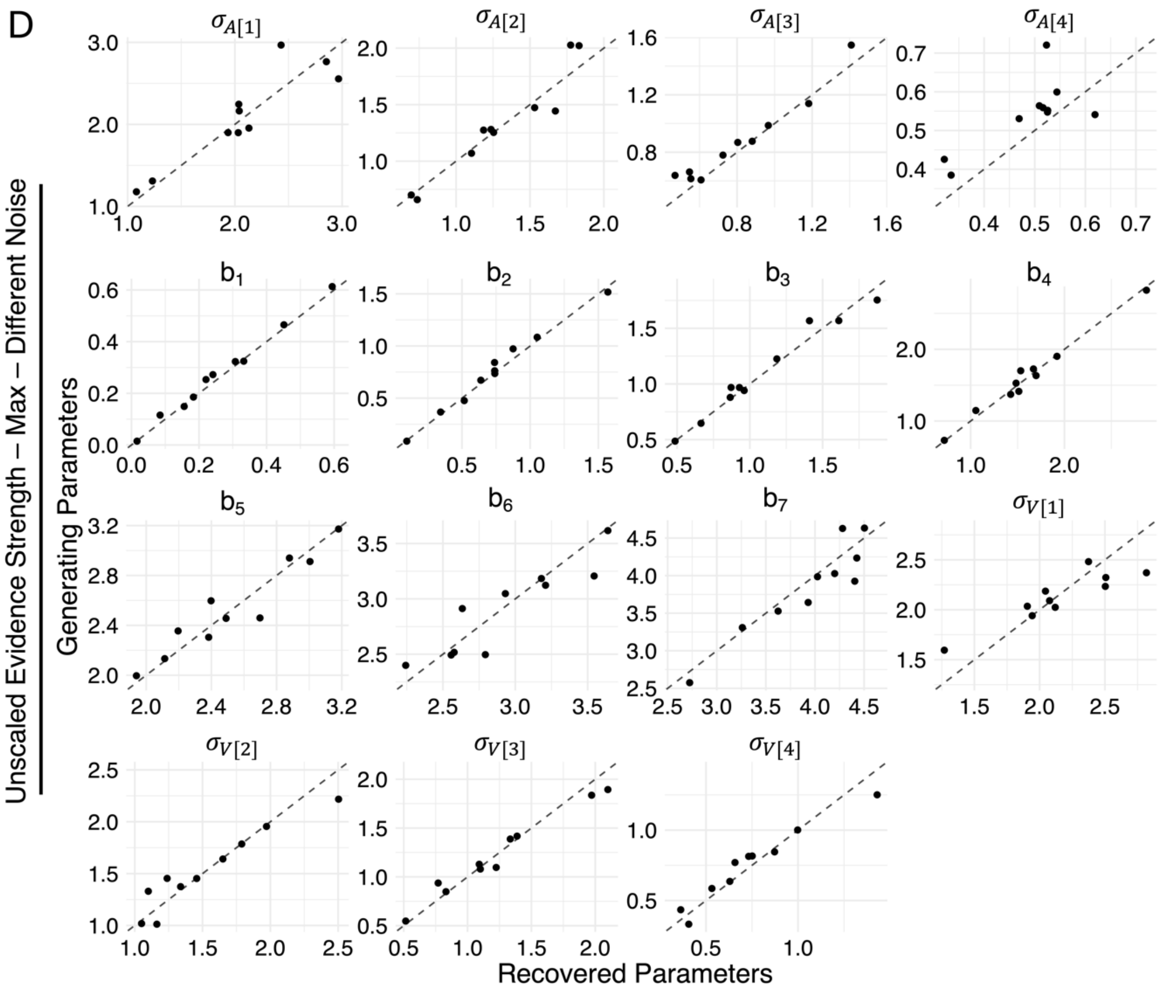


*Note.* Recovery results for (A) integration same noise variant, (B) max same noise variant, (C) integration different noise variant, and **(D)** max different noise variant**.** $b_{1-7}$ refer to the category/confidence boundaries and $\sigma_{1-4}$ refer to the estimated noise parameters at each intensity level (the perceived level of sensory uncertainty in the observer's perception of the stimulus). *w* refers to the weight applied to the visual dimension during integration, where the auditory weight is equal to 1 – *w*. Dotted diagonal lines indicate perfect recovery of the data-generating parameter values.

**Figure S 4**

*Parameter Recovery for Scaled Evidence Strength Models*

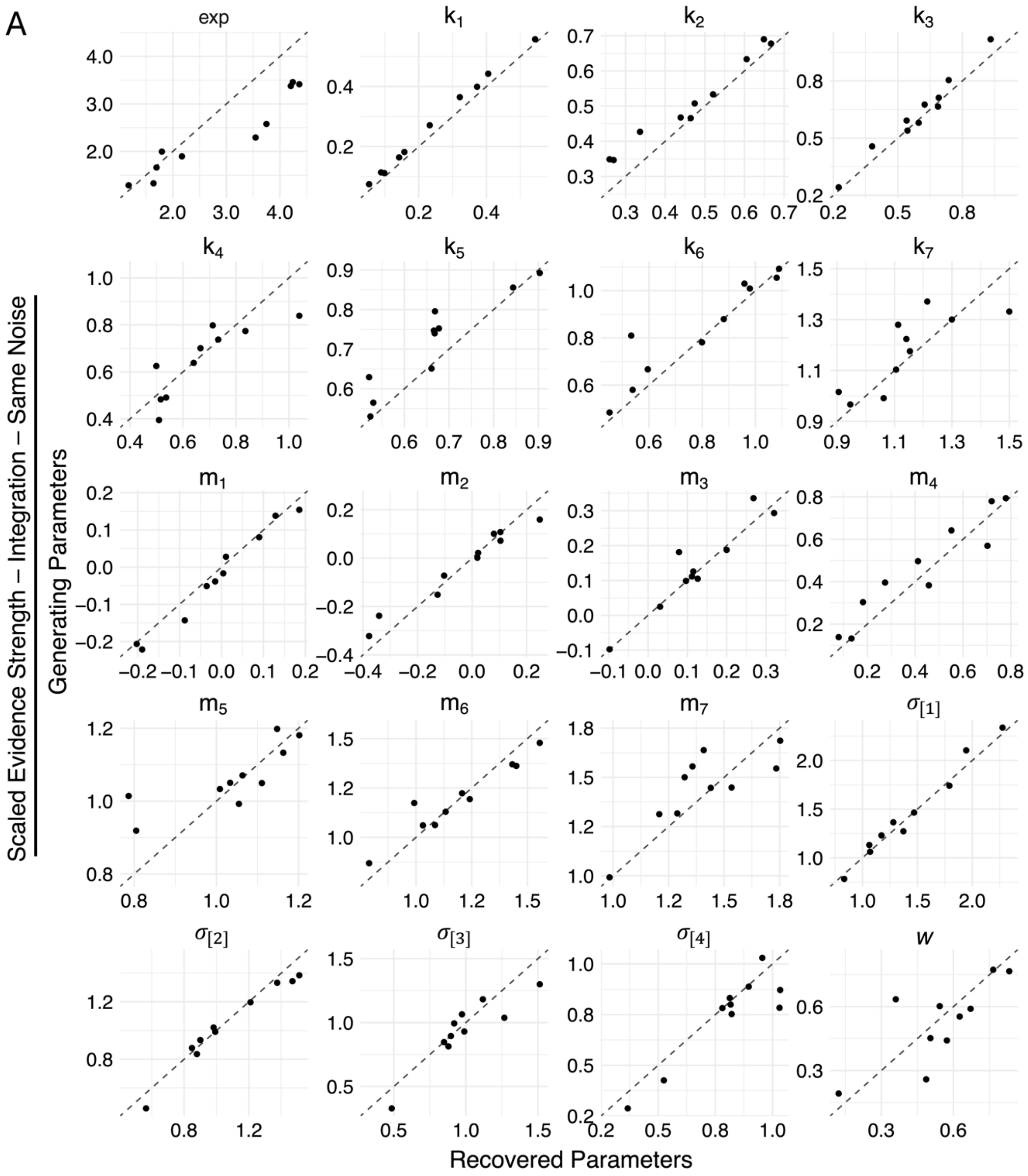


*Note.* Recovery results for **(A)** integration same noise variant, (B) max same noise variant, (C) integration different noise variant, and (D) max different noise variant. $\sigma_{1-4}$ refer to the estimated noise parameters at each intensity level (the perceived level of sensory uncertainty in the observer's perception of the stimulus). Parameters $k_{1-7}$, $m_{1-7}$ and *exp* are used to estimate the category/confidence boundaries according to $b = k + m\sigma^{\exp}$ (see description for scaled evidence strength models for more details). *w* refers to the weight applied to the visual dimension during integration, where the auditory weight is equal to 1 – *w*. Dotted diagonal lines indicate perfect recovery of the data-generating parameter values.

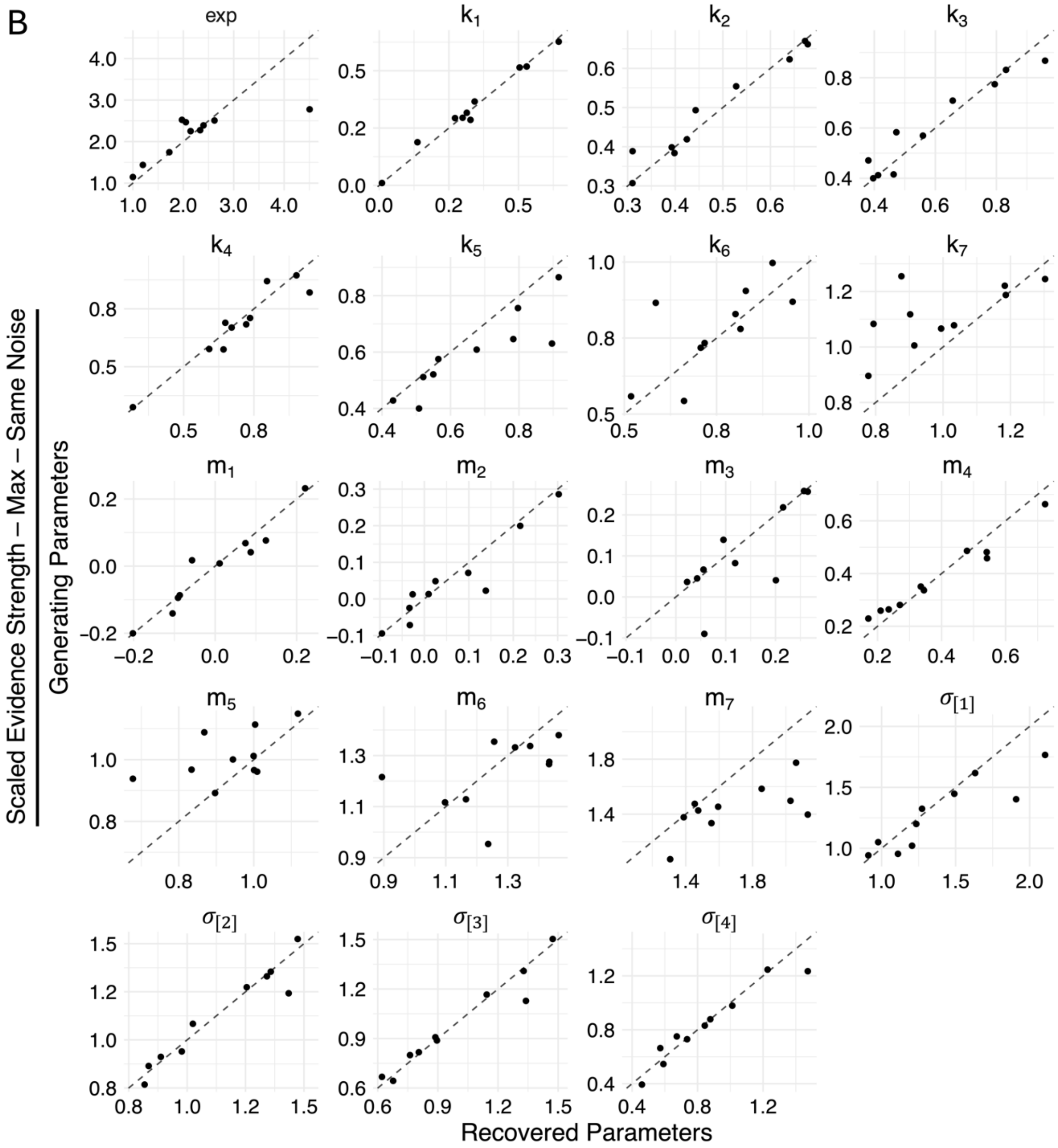


*Note.* Recovery results for (A) integration same noise variant, **(B)** max same noise variant, (C) integration different noise variant, and (D) max different noise variant**.** $\sigma_{1-4}$ refer to the estimated noise parameters at each intensity level (the perceived level of sensory uncertainty in the observer's perception of the stimulus). Parameters $k_{1-7}$, $m_{1-7}$ and *exp* are used to estimate the category/confidence boundaries according to $b = k + m\sigma^{\text{exp}}$ (see description for scaled evidence strength models for more details). *w* refers to the weight applied to the visual dimension during integration, where the auditory weight is equal to 1 – *w*. Dotted diagonal lines indicate perfect recovery of the data-generating parameter values.

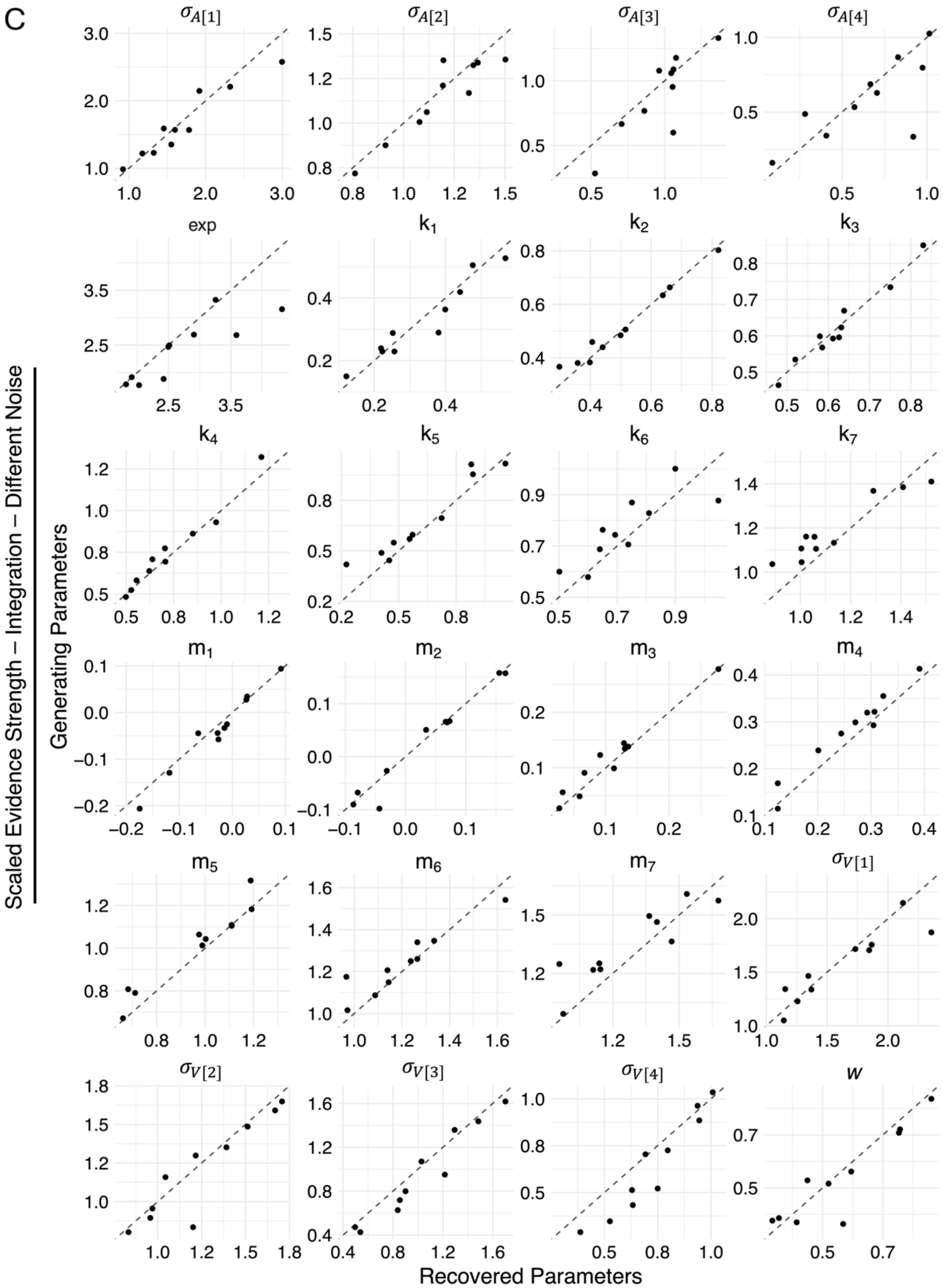


*Note.* Recovery results for (A) integration same noise variant, (B) max same noise variant, **(C)** integration different noise variant, and (D) max different noise variant**.** $\sigma_{1-4}$ refer to the estimated noise parameters at each intensity level (the perceived level of sensory uncertainty in the observer's perception of the stimulus). Parameters $k_{1-7}$, $m_{1-7}$ and *exp* are used to estimate the category/confidence boundaries according to $b = k + m\sigma^{\text{exp}}$ (see description for scaled evidence strength models for more details). *w* refers to the weight applied to the visual dimension during integration, where the auditory weight is equal to 1 – *w*. Dotted diagonal lines indicate perfect recovery of the data-generating parameter values.

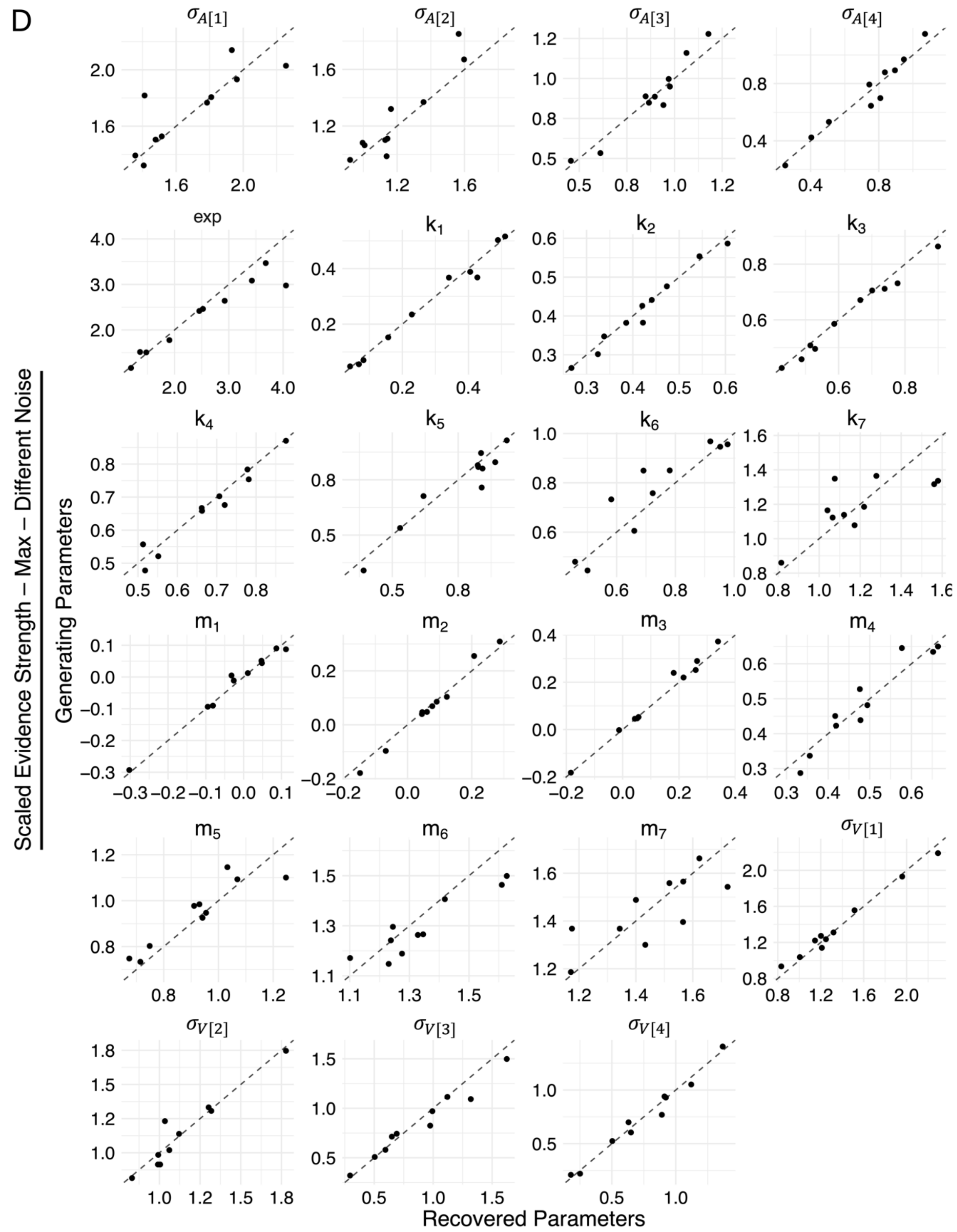


*Note.* Recovery results for (A) integration same noise variant, (B) max same noise variant, (C) integration different noise variant, and **(D)** max different noise variant. $\sigma_{1-4}$ refer to the estimated noise parameters at each intensity level (the perceived level of sensory uncertainty in the observer's perception of the stimulus). Parameters $k_{1-7}$, $m_{1-7}$ and *exp* are used to estimate the category/confidence boundaries according to $b = k + m\sigma^{\text{exp}}$ (see description for scaled evidence strength models for more details). *w* refers to the weight applied to the visual dimension during integration, where the auditory weight is equal to 1 – *w*. Dotted diagonal lines indicate perfect recovery of the data-generating parameter values.

**Figure S 5**

*Parameter Recovery for Bayesian Models*

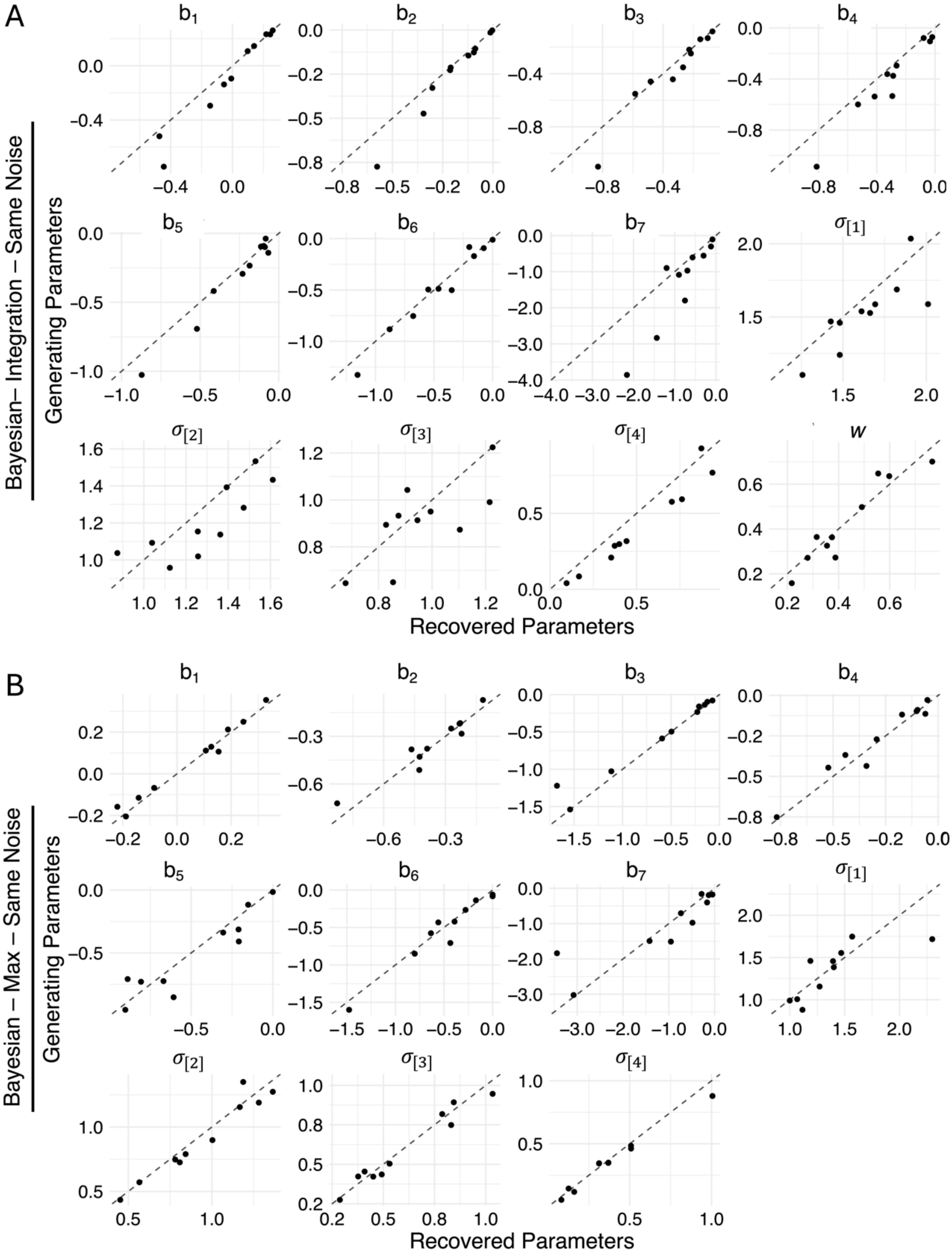


*Note.* Recovery results for **(A)** integration same noise variant, **(B)** max same noise variant, (C) integration different noise variant, and (D) max different noise variant. $b_{1-7}$ refer to the category/confidence boundaries and $\sigma_{1-4}$ refer to the estimated noise parameters at each intensity level (the perceived level of sensory uncertainty in the observer's perception of the stimulus). *w* refers to the weight applied to the visual dimension during integration, where the auditory weight is equal to 1 – *w*. Dotted diagonal lines indicate perfect recovery of the data-generating parameter values.

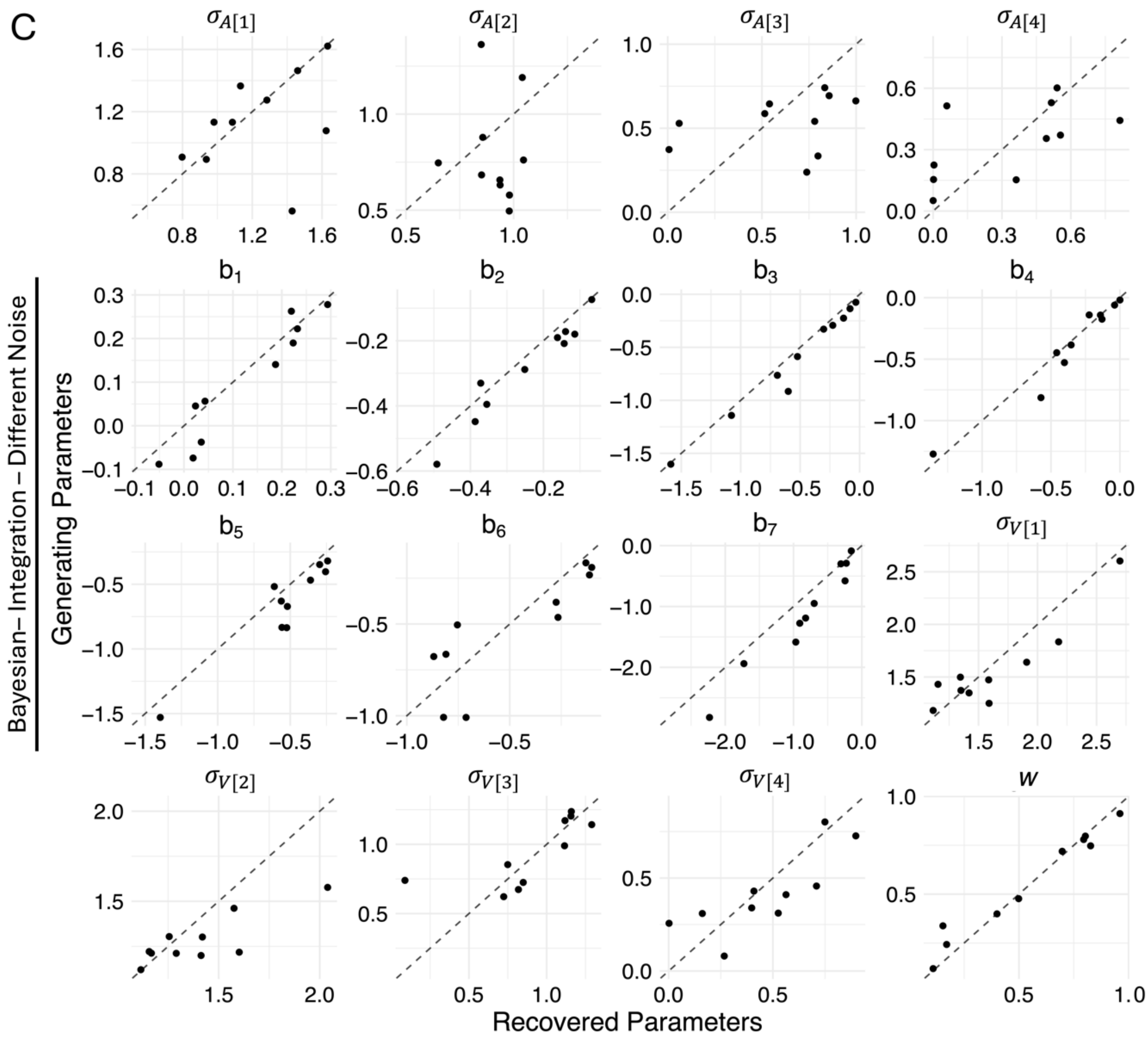


*Note.* Recovery results for (A) integration same noise variant, (B) max same noise variant, **(C)** integration different noise variant, and (D) max different noise variant. $b_{1-7}$ refer to the category/confidence boundaries and $\sigma_{1-4}$ refer to the estimated noise parameters at each intensity level (the perceived level of sensory uncertainty in the observer's perception of the stimulus). *w* refers to the weight applied to the visual dimension during integration, where the auditory weight is equal to 1 – *w*. Dotted diagonal lines indicate perfect recovery of the data-generating parameter values.

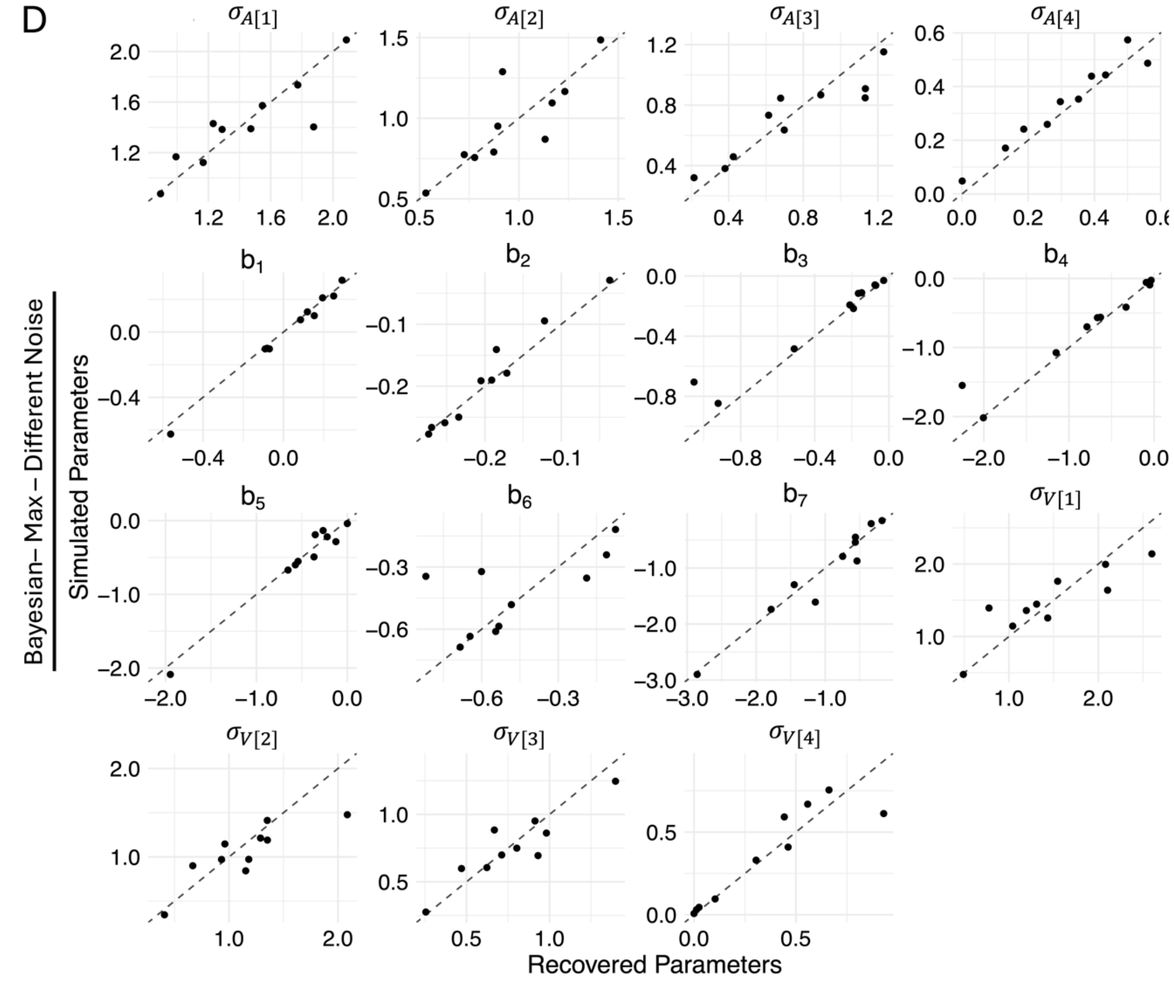


*Note.* Recovery results for (A) integration same noise variant, (B) max same noise variant, (C) integration different noise variant, and **(D)** max different noise variant**.** $b_{1-7}$ refer to the category/confidence boundaries and $\sigma_{1-4}$ refer to the estimated noise parameters at each intensity level (the perceived level of sensory uncertainty in the observer's perception of the stimulus). *w* refers to the weight applied to the visual dimension during integration, where the auditory weight is equal to 1 – *w*. Dotted diagonal lines indicate perfect recovery of the data-generating parameter values.

#### Figure S 6

*Model Recovery*

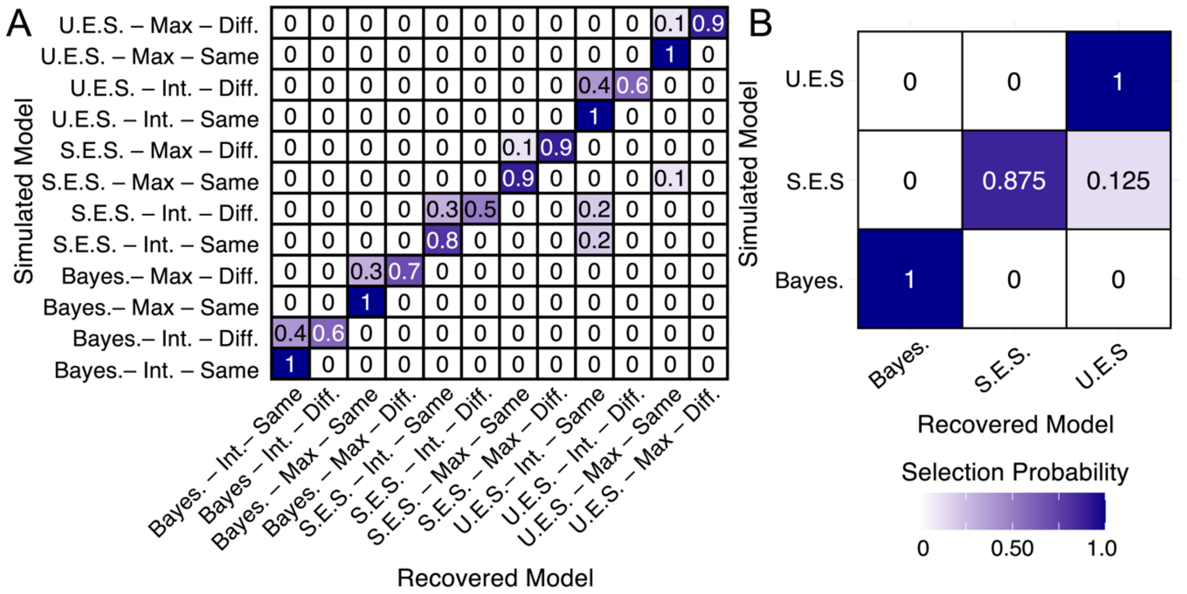


*Note.* Numbers and colours denote the probability that the data generated with model X (y-axis) are best fit by model Y (x-axis). Confusability matrix for **(A)** all model variants and **(B)** model classes using *BIC* for model selection. U.E.S refers to the unscaled evidence strength model class. S.E.S refers to the scaled evidence strength model class. Bayes. refers to the Bayesian model class. Int. refers to the integration assumption. Diff. and same refers to the assumption about the noise parameters across modalities.
